# Effectiveness of Joint Species Distribution Models in the Presence of Imperfect Detection

**DOI:** 10.1101/2021.02.16.431325

**Authors:** Stephanie Hogg, Yan Wang, Lewi Stone

**Affiliations:** Mathematics, School of Science, RMIT, Melbourne, Australia; Biomathematics Unit, School of Zoology, Faculty of Life Science, Tel Aviv University, Israel

**Keywords:** inter-species correlation, detection probability, imperfect detection, joint species distribution model, multivariate probit, occupancy model

## Abstract

1. Joint species distribution models (JSDMs) are a recent development in biogeography and enable the spatial modelling of multiple species and their interactions and dependencies. However, most models do not consider imperfect detection, which can significantly bias estimates. This is one of the first papers to account for imperfect detection when fitting data with JSDMs and to explore the complications that may arise.
2. A multivariate probit JSDM that explicitly accounts for imperfect detection is proposed, and implemented using a Bayesian hierarchical approach. We investigate the performance of the JSDM in the presence of imperfect detection for a range of factors, including varied levels of detection and species occupancy, and varied numbers of survey sites and replications. To understand how effective this JSDM is in practice, we also compare results to those from a JSDM that does not explicitly model detection but instead makes use of “collapsed data”. A case study of owls and gliders in Victoria Australia is also illustrated.
3. Using simulations, we found that the JSDMs explicitly accounting for detection can accurately estimate intrinsic correlation between species with enough survey sites and replications. Reducing the number of survey sites decreases the precision of estimates, while reducing the number of survey replications can lead to biased estimates. For low probabilities of detection, the model may require a large number of survey replications to remove bias from estimates. However, JSDMs not explicitly accounting for detection may have a limited ability to disentangle detection from occupancy, which substantially reduces their ability to accurately infer the species distribution spatially. Our case study showed positive correlation between Sooty Owls and Greater Gliders, despite a low number of survey replications.
4. To avoid biased estimates of inter-species correlations and species distributions, imperfect detection needs to be considered. However, for low probability of detection, the JSDMs explicitly accounting for detection is data hungry. Estimates from such models may still be subject to bias. To overcome the bias, researchers need to carefully design surveys and choose appropriate modelling approaches. The survey design should ensure sufficient survey replications for unbiased inferences on species inter-dependencies and occupancy.

## Introduction

Species distributions models (SDMs) have become an important tool to help manage the environment (Austin 2002; Burgman *et al*. 2005). SDMs model the manner in which the population of a single species depends on environmental covariates such as temperature, water availability, altitude (which are usually known in detail over the entire study area), and may be used to predict the spatial distribution of the population from a set of such covariates. Recently, Joint Species Distribution Models (JSDMs) have been introduced to model multiple species (Ovaskainen *et al*. 2010; Dorazio & Connor 2014; Thorson *et al*. 2015; Harris 2015; Ovaskainen *et al*. 2015). JSDMs may be viewed as extensions of community models which assume that species are distributed independently (Dorazio & Royle 2005). In contrast to community models, JSDMs incorporate statistical dependencies (or correlations) between species that may arise due to species interactions and dependencies, spatial correlations or environmental factors. Some models use hidden environmental factors to account for the statistical dependencies between species (Warton 2015; Hui *et al*. 2015). Other models include a direct intrinsic pairwise correlation between species to account for dependencies (Ovaskainen *et al*. 2010; Dorazio & Connor 2014; Pollock *et al*. 2014).

During ecological surveys not all individuals that are present are observed, a property referred to “imperfect detection”, and one that can create major biases if not corrected. The probability of detection can be influenced by a number of factors, such as observer experience, vegetation cover that interferes with the survey, or difficult weather conditions (Tyre *et al*. 2003). Imperfect detection usually remains, regardless of the expertise of observers. A number of SDMs have attempted to estimate site occupancy probabilities dependent on the degree of imperfect detection (MacKenzie *et al*. 2002; Tyre *et al*. 2003). However, allowance for imperfect detection has previously been incorporated in only several JSDMs (Dorazio & Connor 2014; Rota *et al*. 2016; Tobler *et al*. 2019) and some co-occurrence models for two species (MacKenzie *et al*. 2004; Waddle *et al*. 2010). Apart from the recent paper of Tobler *et al*. (2019), very few papers have thoroughly explored the statistical fitting of JSDM with imperfect detection.

In this paper, we investigate the difficulties of working with JSDMs in the presence of imperfect detection for a range of factors, including varied levels of detection and species occupancy, as well as survey design parameters (i.e. number of sites and survey replications). Using a multivariate probit model (Pollock *et al*. 2014) in conjunction with explicit modelling of detection (Dorazio & Royle 2005), we show that large biases may arise that can lead to erroneous predictions if survey replications are not sufficient, particularly when detection probability is low.

We want to understand how effective the JSDM explicitly accounting for detection is in practice. To do this, we compare models explicitly accounting for detection with models using “collapsed data” across a number of simulated scenarios. Collapsed data summarizes multiple observations or replications into a single measure (Lahoz-Monfort *et al*. 2014). By comparing the models, we determine how well the detection component helps disentangle detection and occupancy, improve parameter estimates, and makes more accurate inferences of the joint species distributions. We find JSDMs which do not explicitly account for detection can be limited in their ability to disentangle detection from occupancy which substantially reduces their ability to accurately infer species distribution.

The paper is organised as follows. We start by introducing a JSDM that explicitly accounts for imperfect detection based on the multivariate probit model developed by Pollock *et al*. (2014). A number of simulated scenarios are explored to evaluate the performance of the JSDM in the presence of imperfect detection under a range of conditions. Simulations have also been carried out to compare outcomes from the JSDM explicitly accounting for detection with those from JSDM that does not explicitly account for detection using collapsed data. The proposed model is then applied to a real case study of Victoria Central Highlands Owls and Gliders. The paper concludes with insights on how well these models estimate inter-species correlations, probabilities of joint occupancy and other parameters in the presence of imperfect detection.

## Materials and Methods

### Modelling occupancy

This paper presents a JSDM that combines occupancy and detection components using a hierarchical approach (Royle & Dorazio 2008; Kéry & Royle 2016). We begin by considering the occupancy component which represents the probability of presence of a species at a typical site, for ease of exposition. A response variable *Z* is used to denote the occupancy status of a single species, with *Z* = 1 if the species is present at a site and 0 otherwise. Species occupancy is modelled using a latent probit approach (Chib & Greenberg 1998; Pollock *et al*. 2014). That is, it is assumed that *Z* is controlled by an underlying unobserved normal latent variable *V* as follows:

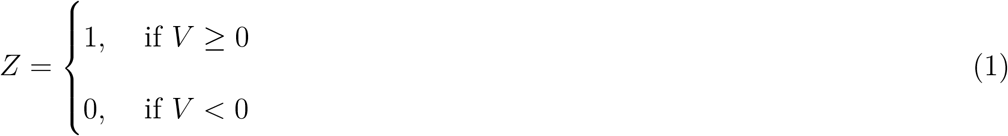

where *V* ∼ *N* (*µ, σ*^2^) and *µ* = *β*_0_ + *X*^*T*^ *β*.

*X*^*T*^ represents the environmental covariates (e.g. temperature or rainfall), and *β* represents the covariate parameters. *µ* is the latent mean which is controlled by latent covariates *X*^*T*^. *σ*^2^ is the intrinsic variance of species’ occupancy. Probit models generate a binary outcome by thresholding a continuous, normally distributed process. Historically, these models were used in toxicology studies where the threshold was considered a tolerance level. Here, the probit models threshold could be interpreted as the point which determines the environmental conditions a species can tolerate or the resources a species needs to survive. This could correspond to the concept of a species niche, where a species has specific resource requirements which need to be met for it to persist (Shoener 2009). Environmental covariates, such as rainfall or vegetation, correspond to the additive resources available at a site.

Extending this model to multiple species at multiple sites, we have *Z*_*ik*_ = 1 if species *k* is present at site *i*, or zero otherwise. Correspondingly, there is a latent variable *V*_*ik*_ associated with *Z*_*ik*_ so that

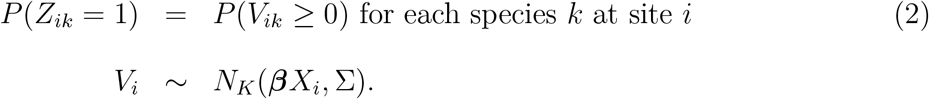

This is the multivariate probit model proposed in Pollock *et al*. (2014). The vector *V*_*i*_ at each site *i*, follows a *K*-dimensional multivariate normal distribution, with the same covariance matrix Σ and mean vector (*µ* = ***β****X*_*i*_). *X*_*i*_ is the vector of environmental covariates values at site *i* and ***β*** is a *K* by *P* coefficient matrix.

To model dependencies between the species, the JSDM uses pair-wise inter-species correlations *ρ*_*V*_ which are defined in equation 2 by the covariance matrix Σ. This matrix represents intrinsic dependencies between the species, such as those arising from species interaction. The diagonal elements Σ_*kk*_ define the intrinsic variance 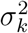 of each species *k*, while off diagonal elements Σ_*kk*′_ define the intrinsic latent covariance *σ*_*kk*′_ between any pair of species *k* and *k*′. The intrinsic latent correlation between any pair of species is calculated as *ρ*_*V*_ = *σ*_*kk*′_ */*(*σ*_*k*_*σ*_*k*′_). The model does not consider higher order correlations between species. *ρ*_*V*_ represents intrinsic correlation only and does not vary with external factors such as the mean occupancy or environmental factors. It is important to realise that *ρ*_*V*_ is not equivalent to the overall latent correlation between species (i.e., correlation between *V*_*k*_ and 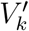), which varies with environmental covariates. Environmental covariates may introduce extrinsic correlation between species, for instance where both species are more likely to occur as average rainfall increases.

Here we give a simple example for two species. Figure 1 is a contour diagram of probability for a bivariate normal distribution. The marginal probability of occupancy of species *k* is denoted by *m*_*k*_ = *P* (*Z*_*k*_ = 1) = *P* (*V*_*k*_ *>* 0). It is calculated as the volume under the curved surface in the right two quadrants for species 1 (*V*_1_ *>* 0) and in the top two quadrants for species 2 (*V*_2_ *>* 0). The probability of co-occurrence, *ω*_11_, is the probability of two species sharing the same site. It is defined as *ω*_11_ = *P* (*Z*_1_ = 1, *Z*_2_ = 1), which corresponds to the volume under the curved surface in the upper right quadrant in Figure 1. Figure 1 shows how the marginal probability of occupancy changes as the means change from a) and b) to c). The marginal probability of occupancy depends on *µ*_*k*_ and is not affected by intrinsic correlation (Tong 1990). However, the probability of co-occurrence changes with both mean vector *µ* and correlation *ρ*_*V*_.

**Figure 1:**
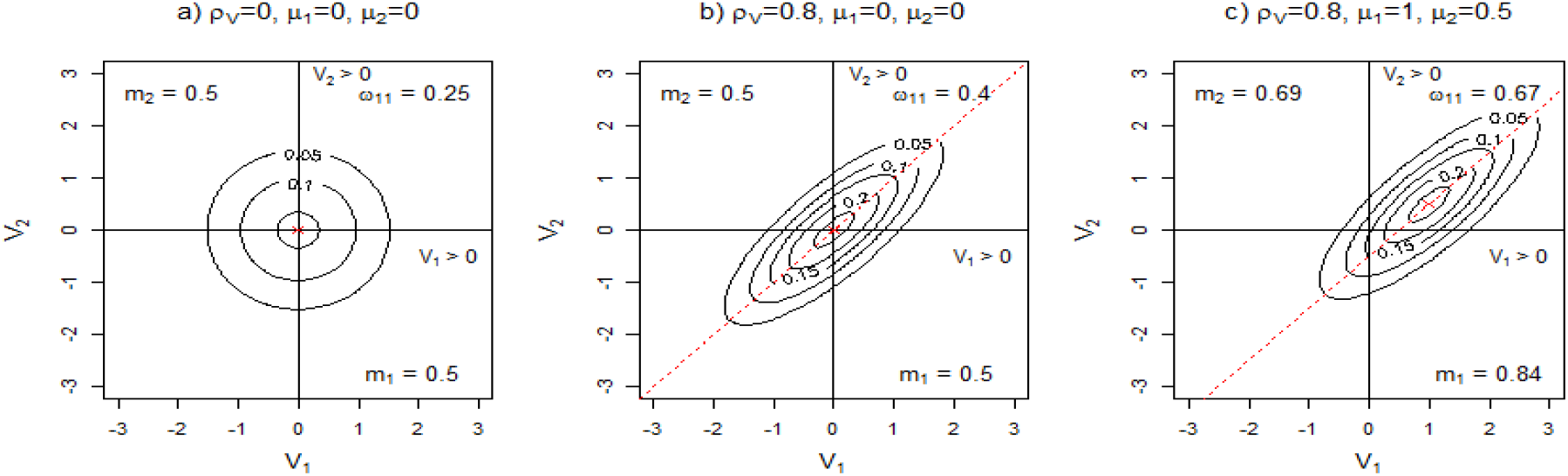
Contour diagram of joint probability of bivariate normal distribution for latent occupancy V_1_ and V_2_: ellipses represent a given joint probability of occupancy. Probabilities are calculated from mean latent occupancy µ_k_ for species k and intrinsic correlation between species ρ_V_. µ_k_ sets the centre of the contour ellipses and determines m_k_, the marginal probability of occurrence. ρ_V_ controls the slope and tightness of the contours. ρ_V_ and µ_k_ determine ω_11_, the joint probability both species occur at the same site.

### Modelling Detection

Due to imperfect detection, we do not always observe the presence of a species at a given site. We define the observed occupancy as *Y*_*ik*_ where *Y*_*ik*_ = 1 if species *k* is detected at site *i* and *Y*_*ik*_ = 0 otherwise. We set the random binary variable *D*_*ik*_ ∼ *Bernoulli*(*p*_*ik*_), where *p*_*ik*_ is the conditional probability of detection for species *k* given that the species is present at site *i*. Then observed occupancy relates to true occupancy (*Z*_*ik*_) as follows (Royle 2004; Mackenzie & Royle 2005; Dorazio & Royle 2005):

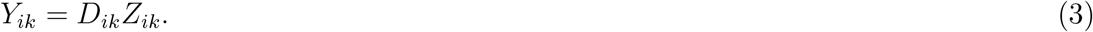

If there is only one replication at each site, the imperfect detection is confounded with the occupancy (Tyre *et al*. 2003; Dorazio 2014). In order to separate detection *D* from occupancy *Z*, our model requires multiple replications of observations from each site. We use *Y*_*ijk*_ to denote the observed occupancy of the *j*^*th*^ replication for species *k* at site *i*.

The ability to detect a species at a site can be influenced by environmental factors, such as wind, terrain ruggedness, or vegetation density, as well as survey factors including time or date of observations, or detection method. When covariates are included using a logit model, the detection model becomes (Dorazio & Royle 2005; Kéry & Royle 2016):

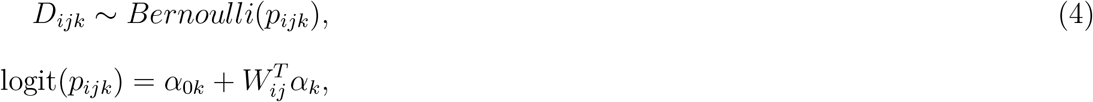

where *W*_*i*_ is the vector of covariates that affect the detection process at site *i, α*_*k*_ is the vector of detection slope parameters and *α*_*k*0_ is the intercept.

### The Complete Hierarchical Model Accounting for Detection

Putting it all together, the full hierarchical model, including both occupancy and detection, for each species *k*, site *i*, and survey replication *j*, is presented below.

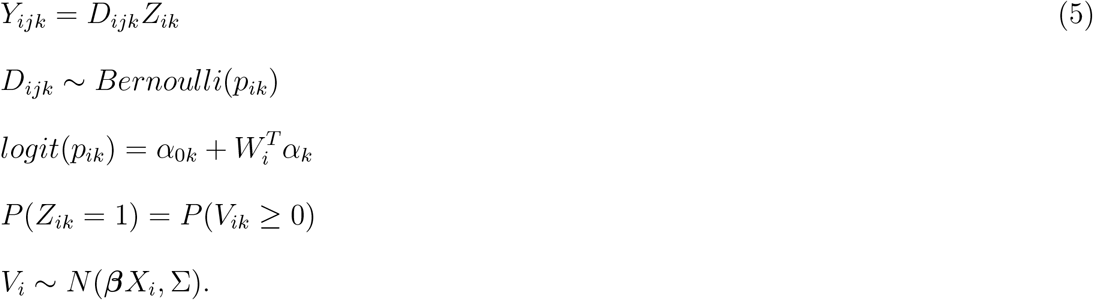

For a multivariate probit model, we require the correlation matrix. The correlation matrix is found by “standardising” the covariance matrix Σ. That is, each element Σ_*kk*′_ is divided by the intrinsic standard deviation of species *k* (*σ*_*k*_) and *k*′ (*σ*_*k*′_), where 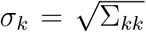. The occupancy parameter estimates also need to be standardized, i.e.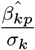, so that they can be interpreted as a standard probit regression coefficient (Chib & Greenberg 1998).

Bayesian analysis has been used to estimate the model following a hierarchical approach. The inverse Wishart distribution is used as the prior for the covariance matrix with a degree of freedom of *K* + 1, where *K* is the number of species. These prior settings follow the practice of Pollock *et al*. (2014) and generally result in a non-informative prior (Hurtado Rúa *et al*. 2015). A sensitivity analysis of covariance priors was also conducted, as described in Supplement S4-2. This analysis compares results for the inverse Wishart priors with two alternative priors (Demirhan & Hamurkaroglu 2008). Estimates of *ρ*_*V*_ do not appear to be sensitive to the prior.

Normal priors were used for occupancy *β* parameters and detection *α* parameters. This is a hierarchical model, so hyperpriors are used to describe the distributions of means and the standard deviations of the normal distributions. The mean hyperpriors were assumed to follow a normal distribution of 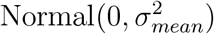, while standard deviation priors followed uniform distributions of Uniform(0, *max*_*sd*_). We investigated estimates of *ρ*_*V*_ and other parameters and their sensitivity to a range of hyperpriors, with the standard deviations (*max*_*sd*_) ranging from 4 to 100 for *β* occupancy priors and from 1 to 100 for *α* detection priors for simulated data. The sensitivity analysis shows no noticeable effect of priors on parameter estimates for simulated data as shown by supplementary Figures S4-3 and S4-2. In later simulations, occupancy hyperpriors used a standard deviation of *max*_*sd*_ ∈ {4, 8} and detection hyperpriors used *max*_*sd*_ ∈ {2, 4}.

We deployed the hierarchical model using the Markov Chain Monte Carlo Bayesian software JAGS (Plummer 2016) run through R 3.5.1 (R Core Team 2015). The JAGS code was based on the work of both Pollock *et al*. (2014) and Kéry & Royle (2016). The code is shown in Supplement S1-3.1 and S1-3.2.

### Simulation Scenarios

We studied four simulation scenarios and a case study of Victorian Central Highland Owls and Gliders. For the former, the occupancy data of the multiple species were simulated following a multivariate probit model (Equation 2) and the detection data was further simulated following a Bernoulli model (Equation 4). We have used two species in our simulation scenarios, but the key results can be generalised to multiple species. For convenience, the correlation matrix is used for the covariance matrix Σ. To investigate the impact of different conditions, multiple independent data sets were simulated for each parameter combination. The parameters of interest were the intrinsic correlation between species (*ρ*_*V*_), occupancy parameters *β* and detection parameters *α*. Two summary statistics, the 95% credible interval and the mean estimate of parameter estimate, are plotted for each of the multiple data sets. We consider two types of errors, precision and bias, to assess model performance. Precision is measured by the width of credible interval, while bias is measured by the distance of the estimated mean to the true parameter values (MacKenzie *et al*. 2002).

#### Scenario 1

We determined if the JSDM accounting for detection could successfully estimate parameters, and the number of sites and survey replications required for accurate estimations. Data was simulated using the occupancy model (Eq 2) with one covariate (*X*_1_), and detection model (Eq 4) with one covariate (*W*_1_). Covariates were generated using a uniform distribution between −1 and 1. Five independent data sets were simulated for each combination of *ρ*_*V*_ ∈ {−0.5, 0, 0.5, 0.9}, *N* ∈ {50, 100, 200, 400}, *R* ∈ {3, 5, 7, 10} and slope parameters (*β*_1*k*_ ∈ {−1, 1} and *α*_1*k*_ ∈ {−1, 0, 1}). The paper shows outcomes for independent covariates with one combination of slope parameters where they are set to one (*β*_1*k*_ = 1, *α*_1*k*_ = 1) for both species. S2-2 shows outcomes for further *α* and *β* parameter combinations for both independent and dependent covariates. Covariates are dependent if an occupancy covariate affects a detection covariate, for instance if type of vegetation is required for occupancy but thickness of vegetation reduces detection.

#### Scenario 2

We investigated how well model performs under a range of possible conditions, such as low probabilities of detection or occurrence, when survey replications may be limited. For this study, species were assumed to have the same, constant probability of occurrence, *m* = *m*_1_ = *m*_2_, and the same, constant probability of detection, *p* = *p*_1_ = *p*_2_. Occupancy data was simulated using Equation 2 where *m* = *φ*(*µ*) = *φ*(*β*_0_), and detection simulated using Equation 4 where *p* = *logit*^−1^(*α*_0_). For each combination of *p* ∈ {0.25, 0.5, 0.75}, *m* ∈ {0.2, 0.5, 0.85}, and *ρ*_*V*_, 14 data sets were simulated for *N* = 200 sites and a number of survey replications, with *R* = 3 shown in this paper. Study details are provided in S2-1 and results for *p* ∈ {0.25, 0.5, 0.75}, *m* = 0.5, *N* = 200 and *R* ∈ {3, 5, 7} are shown in S2-3. Results for the negative correlations *ρ*_*V*_ ∈ {−0.75, −0.5, −0.25} are shown in S2-3.

#### Scenario 3

We are interested in studying how effective the JSDM explicitly accounting for detection is when estimating parameters compared to models which do not explicitly account for detection. For a fair comparison based on the same sampling effort, we used the “collapsed data” approach. This approach provides the most favorable comparison conditions for a model not explicitly accounting for detection (Lahoz-Monfort *et al*. 2014). Data is collapsed to a single record, 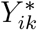 by setting 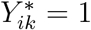 if the species *k* has been detected in any of *R* observations at a site *i*, and 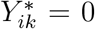 otherwise. Collapsing data as described has the effect of setting the effective conditional detection probability 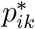 to (MacKenzie *et al*. 2002):

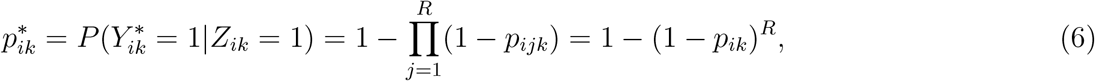

assuming detection probabilities are the same across survey replications. 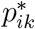 tends to unity 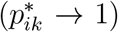 as the number *R* of survey replications becomes large. That is, when R is large the estimate from the collapsed data is close to the true occupancy.

The collapsed data approach effectively has the same information as the explicit model with multiple survey replications, as it uses the same data just in different forms. For example, in terms of the unconditional probability of not detecting a species at a site after *R* surveys, we would obviously have for both models the same probability:

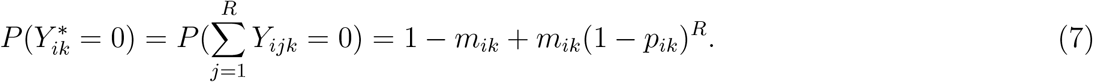

Here *j* is a survey at site *i, R* is the total number of surveys, *m*_*ik*_ is the marginal probability of occurrence, and *p*_*ik*_ is the conditional probability of detection given species *k* occurs at site *i*. Supplement S5-3 shows how the right hand side term in equation 7 is derived.

For this scenario, the simulation data from scenario 1 was collapsed and the occupancy model (Equation 2) fitted to the collapsed data. This model included both occupancy covariates (*X*_*i*_) and detection covariates (*W*_*i*_). We then compared estimates of *ρ*_*V*_, occupancy *β* parameters and detection *α* parameters for the two models.

#### Scenario 4

JSDMs provide insight into joint probability distributions of species which show how species affect each other’s distributions. However, few papers have investigated co-occurrence and the effects of imperfect detection, apart from earlier studies for examining two species co-occurrence (MacKenzie *et al*. 2004; Waddle *et al*. 2010). Recall that *ω*_11_ is the probability two species occur together for any given site. The value of *ω*_11_ is determined by the values of the environmental covariate *X*_1_ and the intrinsic correlation of the species. Estimates of the probability of co-occurrence (*ω*_11_) were evaluated from the results of scenarios 1 and 3 over varying *R* and *N*.

We also evaluated the impact on joint probability of co-occurrence estimates of ignoring intrinsic correlations between species. We compared approximations of *ω*_11_ using single species models with estimates using the JSDM. The joint probability of two species’ co-occurrence can be calculated by using single species models and assuming species have no intrinsic correlation, as follows:

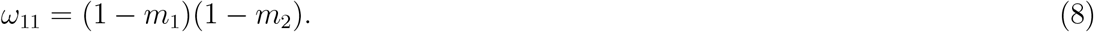

*m*_*k*_ is the marginal probability of occurrence for species *k* as determined by a single species model. We implemented single species species models with detection, using both logit and probit occupancy components as described in Supplement S2-1.1 and fitted them to the scenario 1 data. Joint probabilities were then calculated using Equation 8 and compared to estimates from the JSDM.

## Results

### Scenario 1: Estimation of JSDM parameters for different number of sites *N* and replicates *R*

With adequate replications and site numbers, the JSDM with explicit detection can accurately estimate all model parameters as shown by Figures 2 and 3. However, imperfect detection causes a loss of information in the occupancy data which may badly affect model performance when survey data is insufficient and generate biases.

**Figure 2:**
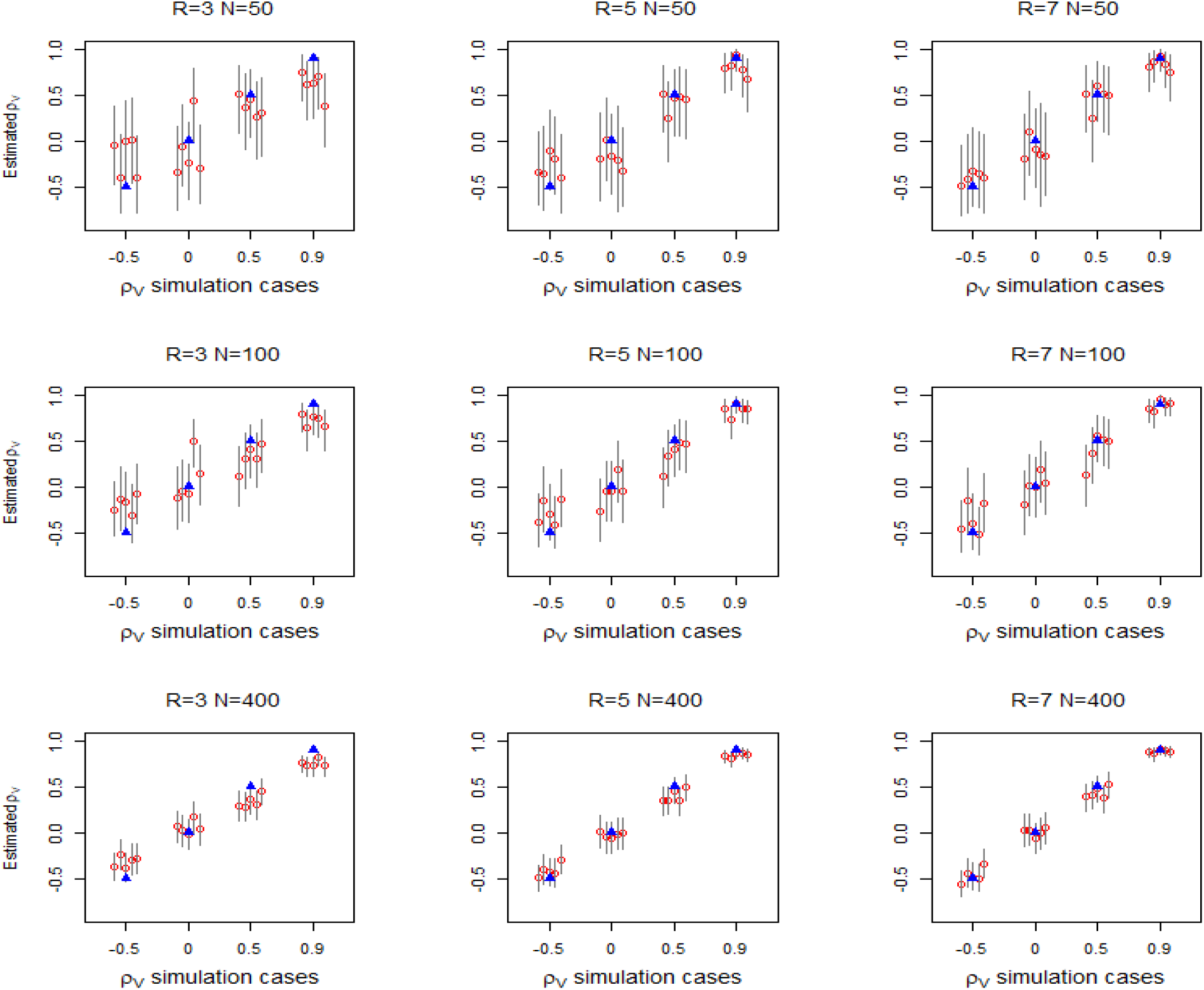
Estimates of *ρ*_*V*_ for different combinations of number of sites *N* and number of replications *R*. Each panel shows the outcome for one combination of *N* and *R* for a number of *ρ*_*V*_ simulation cases. There are 5 simulations per *ρ*_*V*_ case, *ρ*_*V*_ ∈{−0.5, 0, 0.5, 0.9 }. Results are shown as follows: true *ρ*_*V*_ (blue triangle), mean estimate (red circle), and 95% credible intervals limits (grey lines). *N* increases by row, from top (*N* = 50) to bottom (*N* = 400). *R* increases by column from left (3) to right (7).

**Figure 3:**
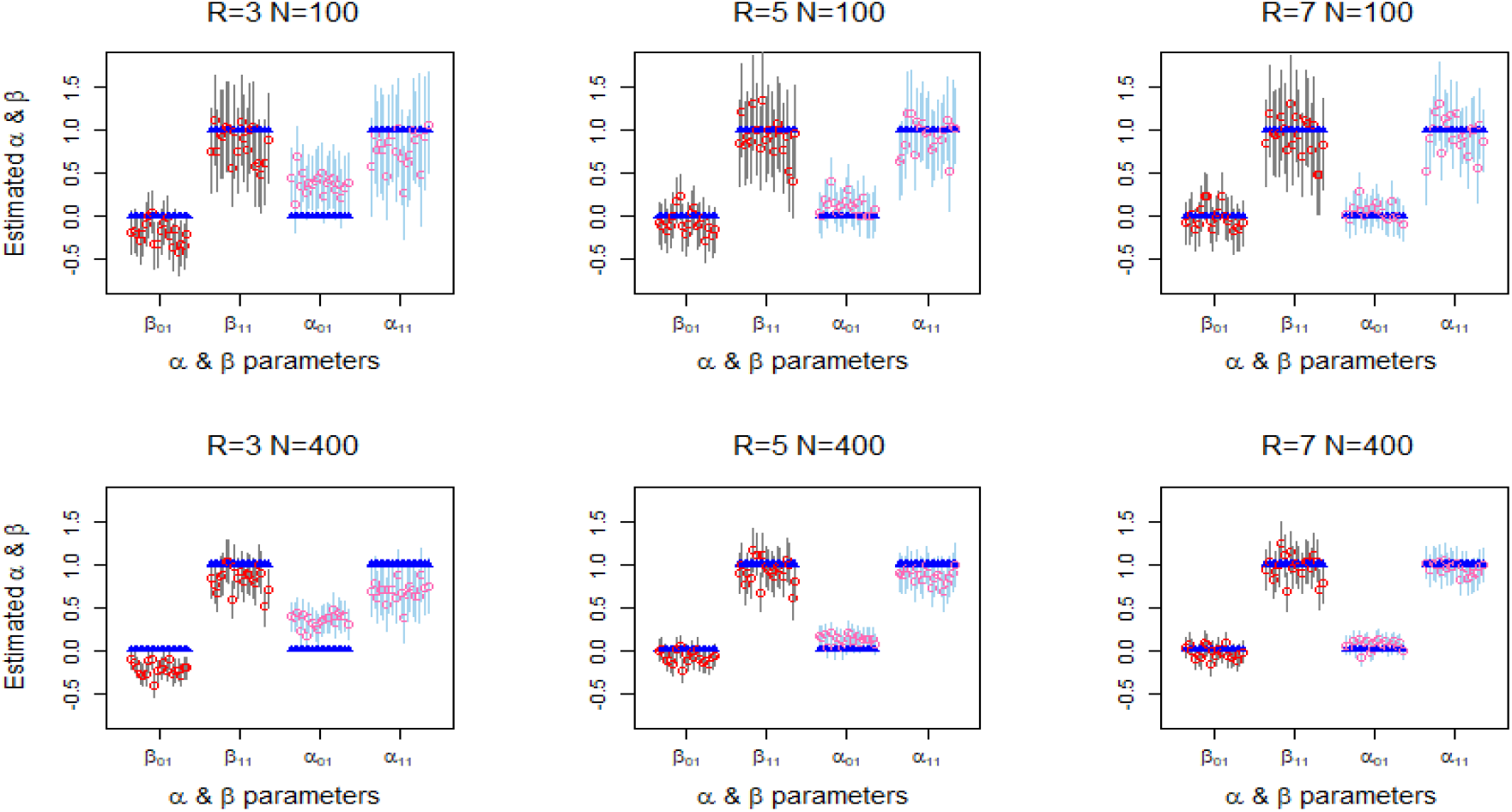
Impact of survey replications *R* on estimates of occupancy and detection parameters. Panels show parameter estimates for *R* increasing from left (3) to right (7) for *N* = 400. Each panel shows 20 estimates for each *β* and *α* parameter as follows: true value (blue triangle), mean estimate (red circle), and 95% credible intervals limits (grey lines) for model accounting for detection.

Figure 2 shows the estimates of *ρ*_*V*_ for different combinations of *N* and *R* for spatially varying occupancy and detection. Estimates of *ρ*_*V*_ become less biased (i.e. mean values get closer to true values) as the number of replications *R* increases, with good estimates for *R* = 7 replications. The estimates show a significant improvement in precision (i.e. the 95% credible interval width of estimates decreased) as the number of sites *N* increases to 400. However, increasing the number of sites *N* does not improve the systematic bias in estimates of *ρ*_*V*_ when controlling for replications.

Figure 3 shows that similar effects of *N* and *R* are evident for estimates of occupancy parameters (*β*) and detection parameters (*α*) in equation 5. When *R* is low (*R* = 3), the magnitude of the covariate coefficients *β*_11_ and *α*_11_ are significantly under-estimated. Figure S2-1 in Supplement S2-2 shows this result extends to both species. As well, it shows occupancy intercepts (*β*_0*k*_) are under-estimated and detection intercepts (*α*_0*k*_) appear to be over-estimated. These biases significantly reduce as the replications increase. Increasing the number of sites *N* did not noticeably improve this bias. However, increasing *N* significantly improved the precision of *β* and *α* parameter estimates, as shown in Figure 3.

Estimates for dependent occupancy and detection covariates are shown in Figure S2-3 Supplement S2-2. For negatively correlated covariates, estimates follow a similar pattern to the independent covariates. For positively correlated covariates, the magnitude of occupancy parameters tended to be overestimated. There was no discernible effect on estimates of intrinsic correlation *ρ*_*V*_.

These results, showing significant bias for low replications (*R* = 3), were consistent across our covariate simulation scenarios, with further examples shown in supplementary Figure S2-2. Figures 2 and Figure 3, indicate that at least *R* = 5 survey replications with *N* = 200 sites are needed to overcome substantial errors in estimates for these simulations where mean probability of detection and occurrence are both *p* = *m* = 0.5. The accuracy of estimates may show further small improvements as number of sites and replications increases further. We also notice that bias (or mean absolute error) is more obvious for higher *ρ*_*V*_, while other conditions are kept the same. More replications are required to reduce absolute errors for 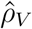, when species are highly correlated.

### Scenario 2: Estimation of JSDM parameters for different mean probabilities of occurrence and detection

Species intrinsic correlation, *ρ*_*V*_, is significantly underestimated when imperfect detection is substantial. The panels in Figure 4 show estimates of species intrinsic correlation *ρ*_*V*_ for different combinations of marginal occupancy *m* and probability of detection *p*. Each row in Figure 4 shows that bias decreases as the probability of detection *p* increases. The third column shows that estimates of intrinsic correlation are reasonably unbiased for high detection probabilities (*p* = 0.75). However, for low or moderate detection probabilities (*p ≤* 0.5), the bias in estimates of *ρ*_*V*_ tends to increase as *m* increases. We also show in supplementary Figure S2-4 that bias in estimating *ρ*_*V*_ decreases as survey replications increase for all values of *p*.

**Figure 4:**
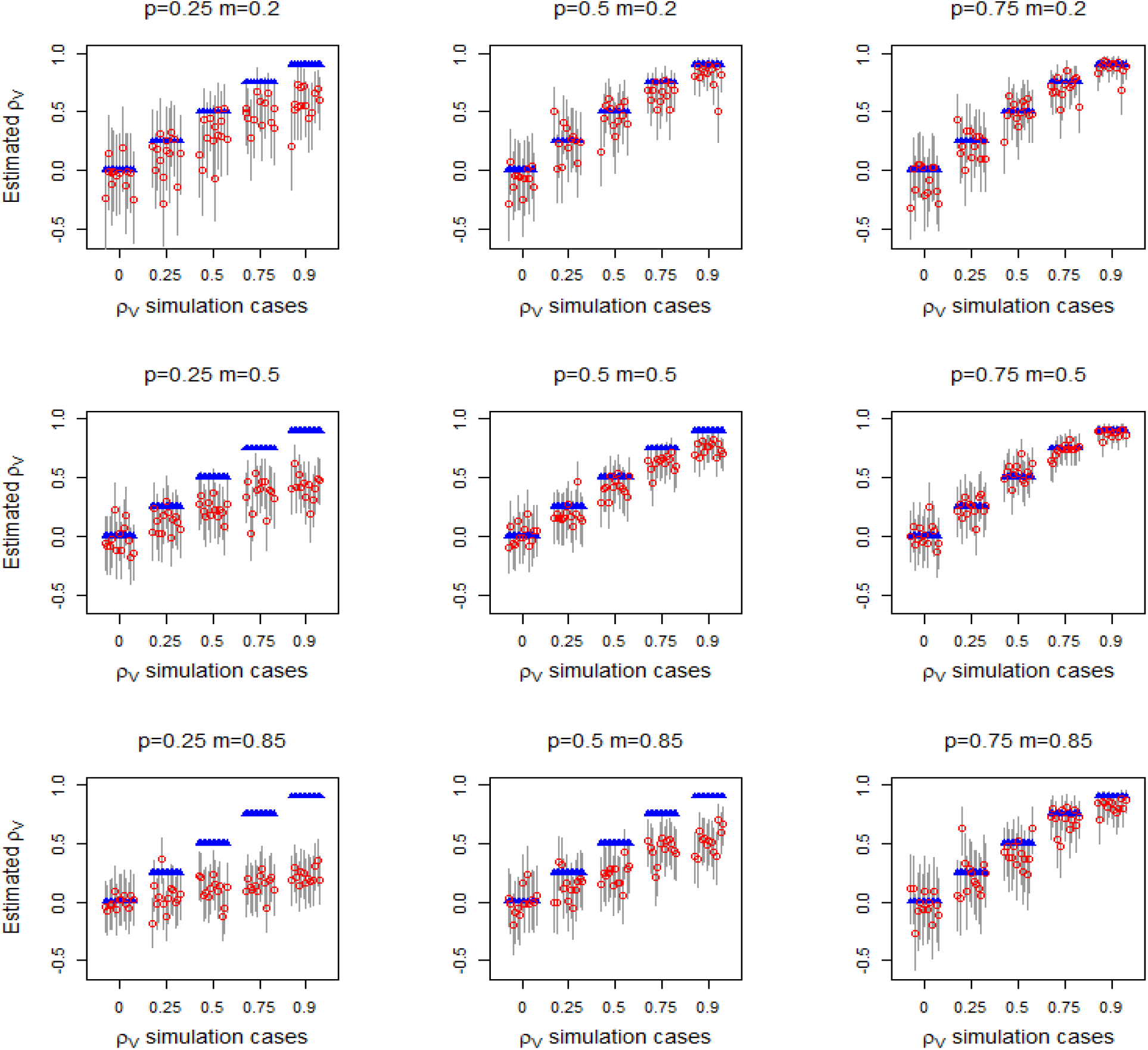
Estimates of *ρ*_*V*_ for different combinations of probability of detection (*p*) and marginal occurrence (*m*) where both species have identical properties. Each panel shows the outcome for one combination of *m* and *p* for a number of *ρ*_*V*_ simulation cases. Here we show results for *ρ*_*V*_ ∈{0, 0.25, 0.5, 0.75, 0.9 } with 13 simulations for each *ρ*_*V*_ case for *N* = 200 sites and *R* = 3 survey replications. The following symbols are used: true *ρ*_*V*_ (blue triangle), mean estimate (red circle), and 95% credible intervals limits (grey lines).

Estimates of negative intrinsic correlation between species were also examined and are shown in supplementary Figure S2-6. Results are similar to estimates of positive intrinsic correlation: sufficient survey replications are required to overcome the errors introduced by imperfect detection.

Figure 5 shows that reasonable estimates of the occupancy and detection intercept parameters (*β* and *α* respectively), can be obtained when the probability of detection is high (*p* = 0.75). When detection probabilities are low to intermediate (*p ≤* 0.5), the *α* detection parameters are overestimated while the *β* occupancy parameters probabilities are under-estimated. The accuracy of estimates of *α* and *β* increase as species detection probabilities increase. Precision also improves as the probability of detection increase.

**Figure 5:**
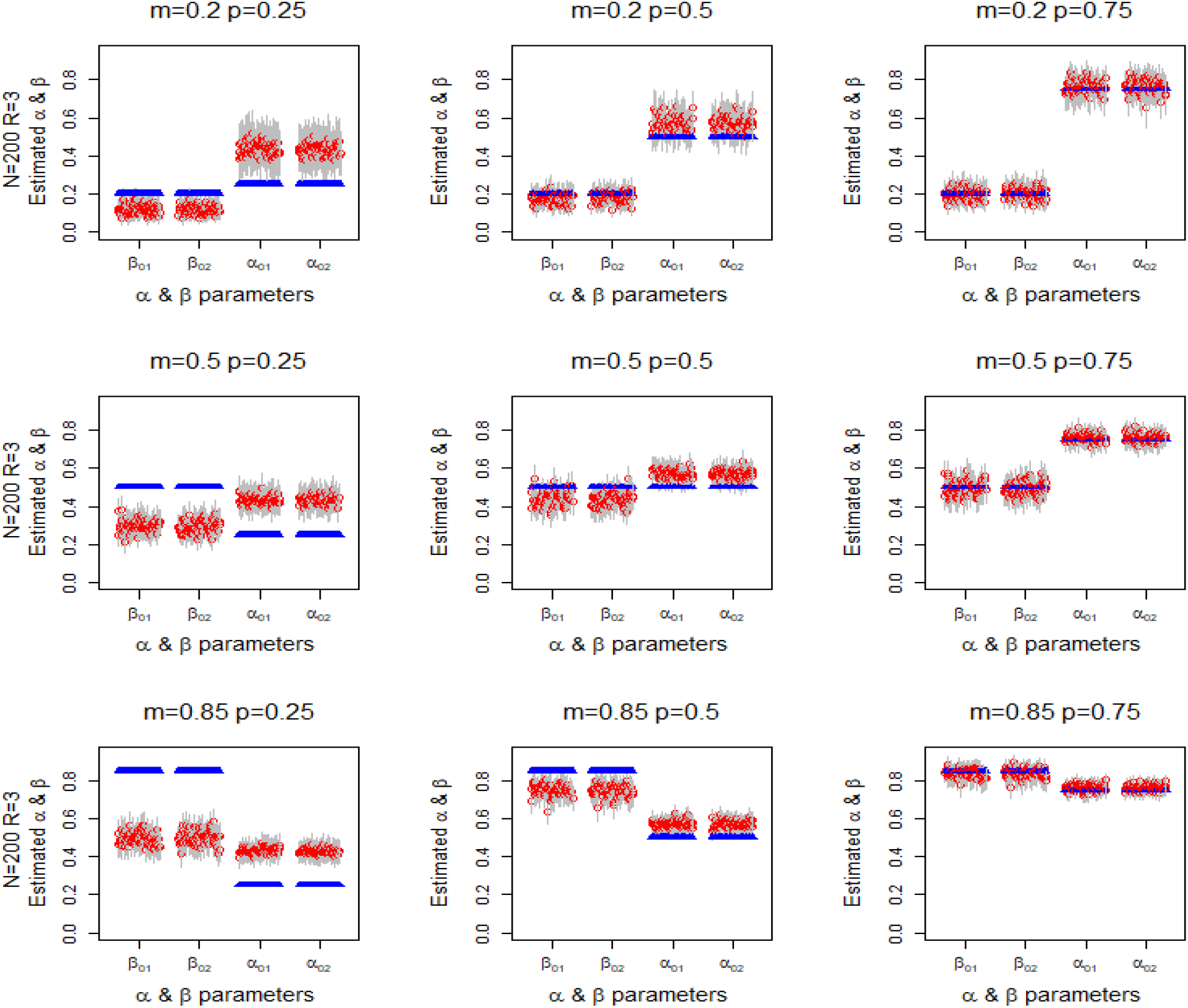
Estimates of occupancy covariate parameters *β* and detection covariate *α* for different combinations of probability of detection (*p*) and marginal occurrence (*m*) where both species have identical properties. Each panel shows the outcome for one combination of *m* and *p* for each *β* and *α* parameter for both species. Here we show results from 65 simulations for each *m* and *p* combination for *N* = 200 sites and *R* = 3 survey replications. The following symbols are used: true parameter value (blue triangle), mean estimate (black circle), and 95% credible intervals limits (red lines).

### Scenario 3: Compare estimates of JSDM parameters between explicit detection and collapsed data approaches

There has been debate on the benefits of explicitly modelling imperfect detection compared to models that do not (Comte & Grenouillet 2013; Guillera-Arroita *et al*. 2014). Figure 6 shows estimates of intrinsic correlation *ρ*_*V*_ for the JSDM models explicitly accounting for detection and for models using collapsed data but not explicitly modelling detection. Estimates of *ρ*_*V*_ were often biased for both models, with both models giving very similar results (in terms of bias and precision) for the same *N* and *R*. Estimates improve in a similar manner as number of sites and replications increase for both models across covariate cases, with further examples in Supplement S2-4. Figure 6 and Figure 2 show equally good estimates of *ρ*_*V*_ for 400 sites and 7 survey replicates for both models.

**Figure 6:**
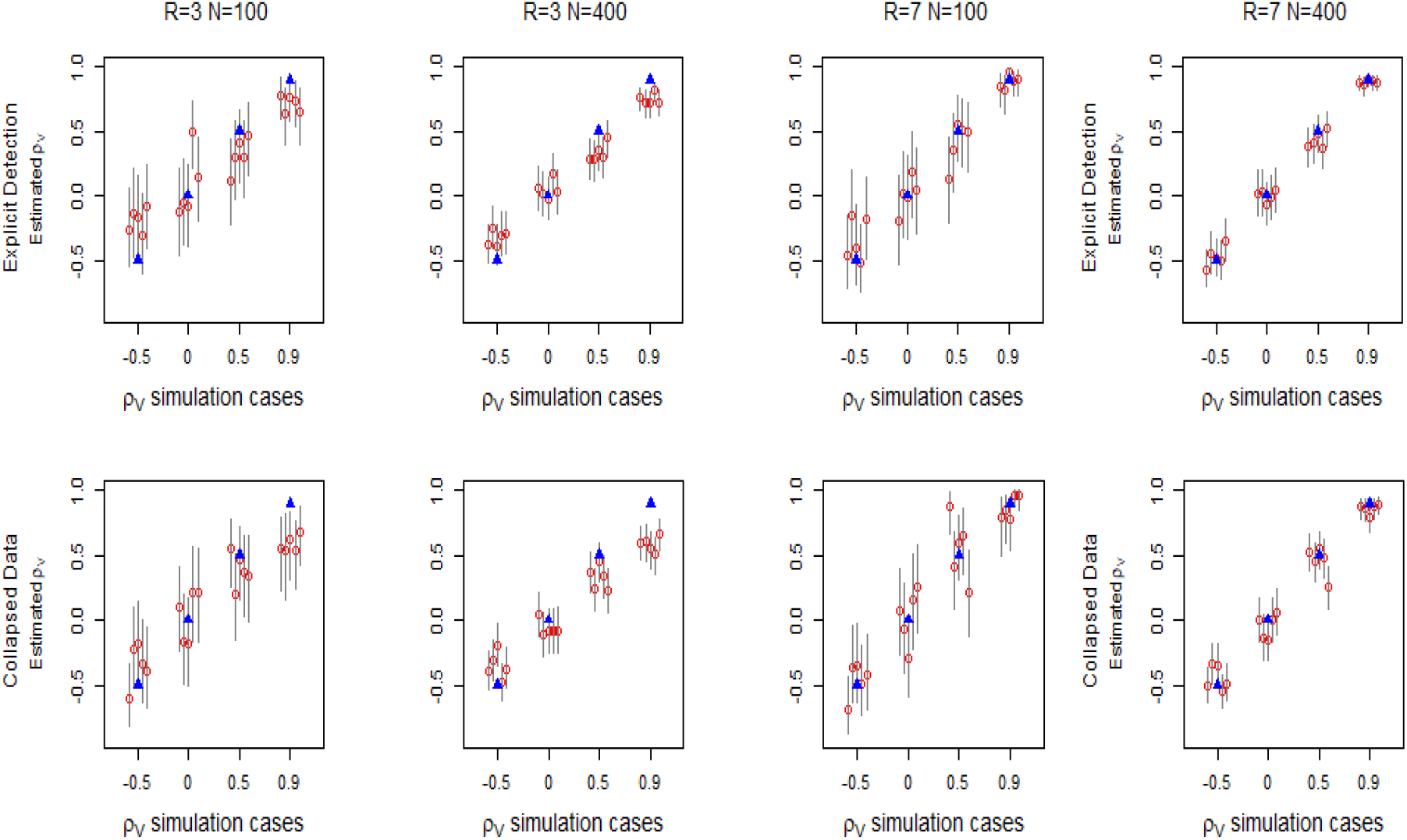
Impact of using collapsed data to estimate *ρ*_*V*_ for varying number of sites *N* and survey replications *R*. Top row shows estimates for the model accounting for detection. Bottom row shows estimates for the model using collapsed data for equivalent *N* and *R*. Each panel shows the outcome for one combination of *N* and *R* for four *ρ*_*V*_ simulation cases. There are 5 simulations per *ρ*_*V*_ case, *ρ*_*V*_ ∈{−0.5, 0, 0.5, 0.9 }. Results are shown as follows: true *ρ*_*V*_ (blue triangle), mean estimate (red circle), and 95% credible intervals (grey lines).

We also compared estimates of coefficients *α* and *β* when detection is modelled explicitly versus the collapsed data approach, as shown by Figure 7. As for scenario 1, increasing *N* improves the precision of parameter estimates for both approaches. Estimates of *β*_1*k*_, the coefficients for environmental covariate *X*_1_, are similar for both models, with bias decreasing as replications *R* increase. Figure 7 shows good estimates of *β*_11_ parameter for both models for 400 sites and 7 survey replicates. For the explicit detection model, bias in estimates of *α*_11_, the parameter for detection covariate *W*_1_, also decreases as *R* increases, with good estimates achieved with seven replications. However, for the model using collapsed data, estimates of *β*_21_, the parameter for *W*_1_, become closer to zero as *R* increases. This is expected as the effective probability of detection for the collapsed data becomes closer to one as the number of replicates increases.

**Figure 7:**
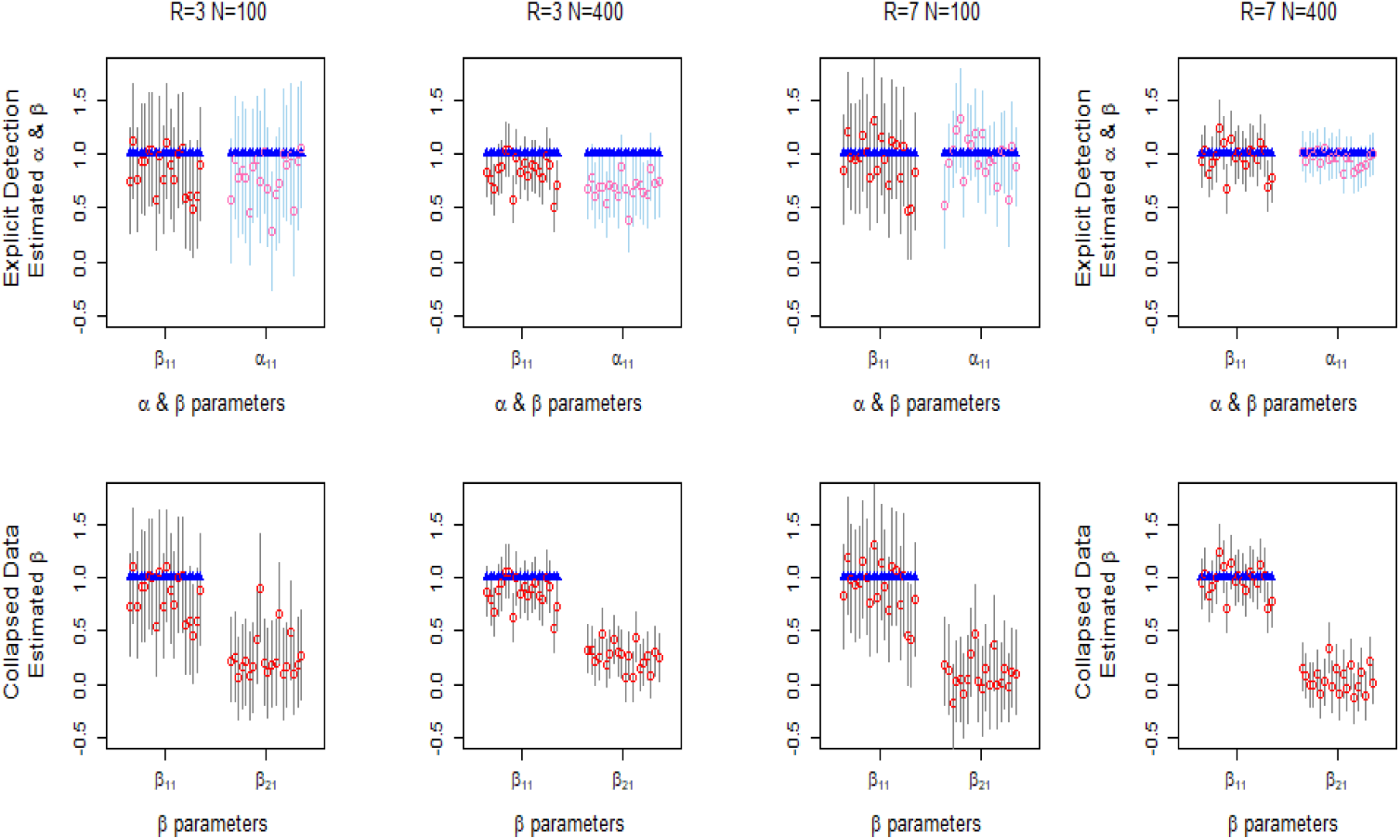
Impact of using collapsed data to estimate occupancy parameters for varying number of sites *N* and survey replications *R*. Top row shows estimates for the model accounting for detection. Bottom row shows estimates for the model using collapsed data for equivalent *N* and *R*. Each panel shows 20 estimates of each *β* or *α* parameter as follows: true value (blue triangle), mean estimate (red/pink circles), and 95% credible intervals (grey/blue lines).

### Scenario 4: Compare estimates of joint probability of occupancy for different modelling approaches

Unlike single species SDMs, JSDMs can determine the joint probability of species co-occurrence across a geographic area. Figure 8 compares actual and predicted values of *ω*_11_ for the JSDM and collapsed data approaches. The figure shows that the true value of *ω*_11_ (left column) increases as *X*_1_ increases, but it is not affected by the detection covariate *W*_1_, as shown by the vertical pattern. Estimates of *ω*_11_ by the model explicitly accounting for detection (centre column), show a similar pattern to the true *ω*_11_ values. The model does tend to under-estimate *ω*_11_, although estimates improve as the number of replications and as the number of sites increases. This result is consistent across covariate combinations in these simulation scenarios, as shown in Supplement S2-5.

**Figure 8:**
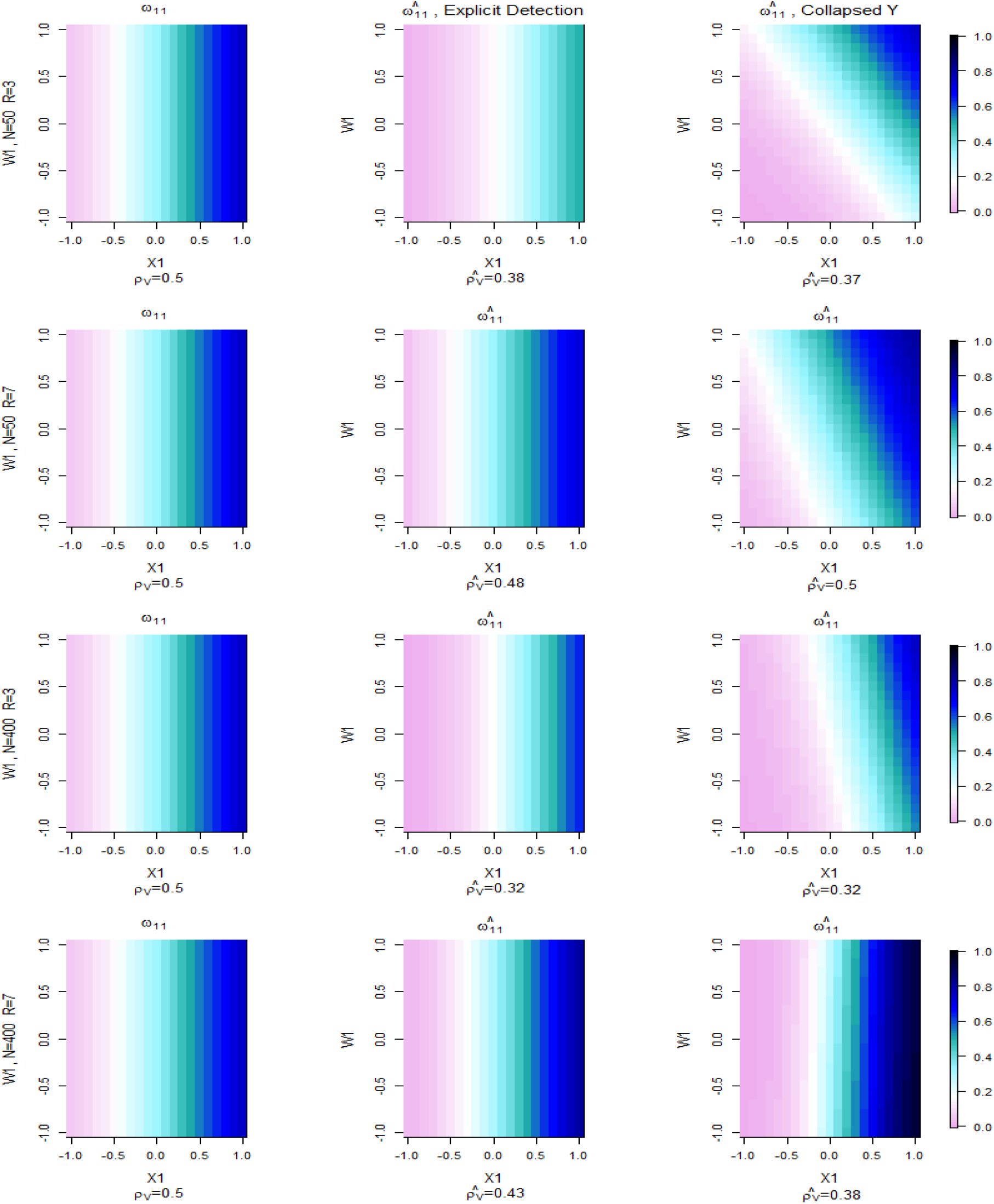
Comparison of true value of *ω*_11_ (left) with predicted values of 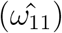 for model explicitly accounting for detection (middle) and model using collapsed data (right). Each plot shows colour coded values of *ω*_11_ (or 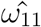) versus covariates *X*_1_ and *W*_1_. Each row shows the results for different combinations of number of sites (N) and survey replications (R).

Estimated values of *ω*_11_ for the collapsed data model are shown in the right-hand column in Figure 8. The fit is poor, especially for low number of sites *N ≤* 100 and replications *R* = 3, with dependence on the detection covariate *W*_1_ evident. The dependence on *W*_1_ is an artefact of including the detection covariates in the occupancy model. However removing these covariates could result in greater errors in estimating occupancy and correlation parameters (Lahoz-Monfort *et al*. 2014). This result for collapsed data models was consistent across the covariate combinations we studied, with further examples shown in Supplement S2-5.

We also estimated an approximation for the joint occupancy *ω*_11_ using single species models, which assume no intrinsic correlation between species. This required estimating the marginal probabilities and inserting them in Equation 8. Figure S2-12 in Supplement S2-6 shows estimates of the joint probability of occupancy for these single species models are not as accurate as the multivariate probit JSDM, when intrinsic correlation is non-zero and number of survey replications is adequate. Estimates of the joint probability of occupancy by single species models are biased, regardless of the number of survey replications if the intrinsic correlation between species is significant. Further details can be found in Supplement S2-6.

### Victorian Central Highland Owl and Glider Study

To illustrate the explicit detection and collapsed data approaches, we applied the methods to presence-absence data from a 2012 study of the Powerful Owl *(Ninox strenua)*, the Sooty Owl *(Tyto tenebricosa)*, the Greater Glider *(Petauroides volans)* and the Yellow Bellied Glider *(Petaurus australis)* in Victoria, Australia. This data was collected over 202 sites in the Central Highland area of Gippsland in Victoria, using call playback and spotlighting with a visual search along a 100m transect, by teams from the former Victorian Department of Environment, Land, Water and Planning (DEPI) (Lumsden *et al*. 2013). Surveys took place from March to August 2012 with two to three survey replications at each site. The targeted species are known to have various interactions, including predator-prey relations and competition for nesting sites (Kavanagh & Bamkin 1995; Kavanagh 2002). Both GIS and site specific covariate data were available and used in our analysis. We selected covariates based on previous simulation scenarios (Lumsden *et al*. 2013), with further variable selection during model fitting. For occupancy modelling we considered eight covariates including seasonal rainfall, elevation, terrain type and vegetation indices. For detection, we considered wind speed, season (autumn or winter), and vegetation type. As the same detection method was used for all surveys, no detection method covariate was included.

First, we fitted a model with explicit detection to the data, using two thirds of the sites as a training set and the other third as validation data. Estimates of occupancy parameters in Table S3-2 show that Powerful Owls, Sooty Owls, and Yellow Bellied glides had a significant negative correlation with elevation, which accords with previous findings (Lumsden *et al*. 2013). Greater Gliders had a significant positive correlation with with NVDI, an indicator of vegetation lushness, in line with previous simulation scenarios (Lumsden *et al*. 2013). They also had a positive correlation with gullies, as shown by the significant negative correlation with Topographic Position Index (TPI). Previous studies found the occupancy of the Greater Gliders correlate with wet, rugged terrain, so TPI may be acting as a proxy for these covariates (Wintle *et al*. 2005; Lumsden *et al*. 2013). Deviance information criterion (DIC), a Bayesian analogue of BIC, was used to select other covariates for inclusion in the model. For species detection, as shown by Table S3-3, both owl species had negative correlation with wind speed. While previous studies have found detection correlated with season for both glider species, no significant effect was found here.

Table 1 shows the latent, intrinsic correlation estimates between species pairs when fitting the explicit model (shown in the *explicit* columns). It shows one positive significant intrinsic correlation between Sooty Owls and Greater Gliders. Sooty Owls are opportunistic predators of Greater Gliders, so this could cause attraction, and hence correlation, between the species. There are indications of positive intrinsic correlation between Powerful Owls and Sooty owls and negative intrinsic correlation between Greater Gliders and Yellow Bellied Gliders, but neither is significant. Credible intervals for correlation are relatively large, so finding significant low-level correlation is difficult. We also fitted models with two and three species, but estimates for *β* occupancy parameters and intrinsic correlation *ρ*_*V*_ are not significantly different from those for the four species model.

**Table 1:**
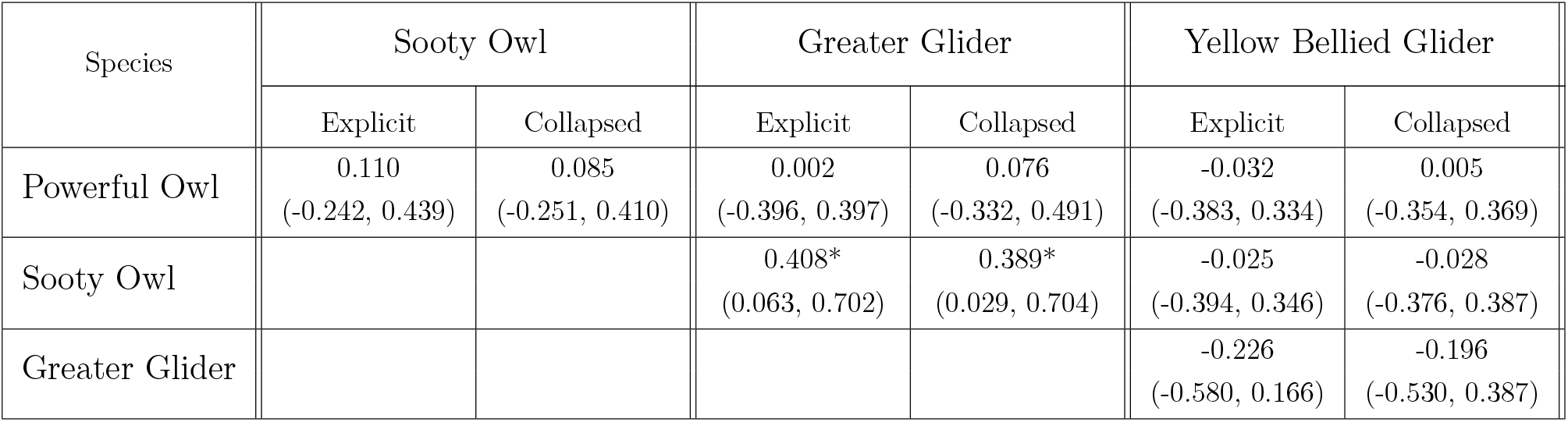
Intrinsic Correlation Estimates for Owl and Glider Data For Models with Explicit Detection and Models using collapsed data. Table shows mean estimated intrinsic correlation values on with 95% credible interval values in brackets below. Results are shown for each distinct pair of species, where second species differs from the first species.

The model fitting was repeated for a single survey replication with no allowance for detection. The correlation between Sooty Owls and Greater Gliders was found to be no longer significant, as shown by Table S3-5. Nor were any other correlations significant. So inter-species dependencies were completely missed when imperfect detection was not taken into account.

We also compared prediction ability for the JSDM explicitly accounting for detection with a JSDM using collapsed data. Predictions were compared by calculating the Area Under the Receiver Operating Curve (AUC) value for each model based on the validation data set. Table S3-4 shows parameter estimates for the collapsed data approach. The detection covariates from the detection component of the JSDM explicitly modelling detection are included in the occupancy model for the collapsed data approach. Supplement 3 Table S3-6 shows these models a have similar fit based on the AUC measure, with neither having a marked advantage over the other. This outcome is in line with previous studies which show that models explicitly accounting for detection and models using collapsed data but not explicitly accounting for detection show similar fit based on AUC measures (Guillera-Arroita *et al*. 2011; Lahoz-Monfort *et al*. 2014). The JSDM with explicit detection can also estimate true probability of occupancy and detection, but these values cannot be verified.

Table 1 compares intrinsic correlation results for the two models. In general, correlation estimates are similar, with both models finding a significant positive correlation between Sooty Owls and Greater Gliders. Credible intervals of all parameters were also large for both models. The collapsed data model uses the wind speed covariate which is observed at sites during the survey. Wind speed data is not available across the area, which limits this model’s ability to make predictions beyond specific sites. Removing wind speed from the model could bias other parameter estimates. This is indicative of the problems that arise when using collapsed data models.

## Discussion

This paper is one of the first studies to investigate the performance of JSDMs in the presence of imperfect detection. This has been carried out through the investigation of extensive simulations over a wide range of combinations that are associated with varied levels of inter-species correlation, different species’ responses to the environment, and effects of spatially varying detection probability, as well as numbers of sites and replications. We have found the JSDM fitting procedure retrieves good estimates of all parameters and true occupancy estimates when the number of survey replicates was sufficiently high. However, if there are insufficient replicates, the model does not adequately allow for imperfect detection, and many of the parameter estimates are found to have surprisingly large levels of bias. Previous studies of single species SDMs with imperfect detection have also reported bias in parameter estimates (MacKenzie *et al*. 2002; Wintle *et al*. 2004; Kéry & Royle 2016). This study shows bias persists with JSDMs and can result in significant underestimation of species intrinsic correlations *ρ*_*V*_. Under some detection conditions, a high number of replications is required for accurate and precise estimates, which may not be feasible for realistic ecological studies, especially if the survey effort available is constrained by the costs and resources required.

To ensure the above mentioned bias was not the product of the priors, a sensitivity analysis was conducted with a series of uniform priors for the Σ covariance matrix (Demirhan & Hamurkaroglu 2008). The sensitivity analysis produced similar estimates of *ρ*_*V*_ to the Inverse Wishart prior. We also investigated the effect of changing the normal hyperpriors for the detection and occupancy covariates, by varying the upper bound of the standard deviation using a uniform distribution. These upper bounds allowed us to see the effect of a non-informative hyperprior (upper bound of 100), to potentially informative hyperpriors (upper bound of 1) for 200 or 400 sites. The varying priors have no significant impact on the posterior distributions of *ρ*_*V*_ or other parameters, as shown in supplementary Figure 2. Moreover, the priors should be considered as weakly or largely non-informative for the restricted (-1,1) range of the simulation parameters (Lemoine 2019). Further, Diaconis & Freedman (1986) formally prove, under certain regularity conditions, that the impact of the prior declines as the sample size increases. Our sensitivity analysis was carried out with 200 sampling sites, which may also explain the lack of effect of the priors. Hence we can conclude our results are influenced by the likelihood rather than the priors for these studies. In practice, it is not uncommon to survey a large number of sampling sites, such as N=202 in our case study, so we would expect these results to apply to realistic studies.

Previous studies of survey effort use asymptotic variance of the occupancy estimator (Fisher information) to estimate the survey effort required for single species models (Mackenzie & Royle 2005; Guillera-Arroita *et al*. 2010). These studies show the trade offs between replication *R* and site numbers *N* required to reach acceptable levels of accuracy. Here, we use Bayesian estimates for simulated data to provide insight to the effort required to estimate correlation accurately. Increasing the number of sites *N* and survey replications *R* both improve estimates, but they behave differently. Figures 2 and 3 show that increasing *N* improves precision (i.e. reduces the 95% credible interval width) of all parameter estimates including intrinsic correlation *ρ*_*V*_. Figures S2-1 shows that precision improves markedly when *N* changes between 100 and 200 sites, with a further relatively minor improvement from 200 to 400 sites. These figures also show that increasing *R* reduces the bias in parameter estimates. Low replications *R* may result in the significant underestimates of the magnitude of of *ρ*_*V*_, occupancy (*β*) parameters and detection (*α*) slope parameters. This bias is not be reduced by increasing *N*. The impact of *R* on precision is minor and does not compensate for an overly low number of sites *N*.

Our simulation results indicate the number of survey replications required to reduce bias varies with the probability of detection *p* and the marginal probability of occupancy *m*. Species with a high probability of detection (*p* = 0.75) may estimate the intrinsic correlation (*ρ*_*V*_) with little bias for *R* = 3 survey replications as shown in Figure 4, yet species with a moderate probability of detection (*p* = 0.5) may require at least *R* = 5 survey replications as shown in Figures 2. Figure S2-4 shows species with a low probability of detection (*p* = 0.25) may require *R* = 7 or more survey replications. These results may be further affected by underlying marginal occupancy *m* if the probability of detection is low. Figure S2-4 shows that the number of survey replications required to estimate *ρ* adequately increases with *m* when *p ≤* 0.5. We also found increases in absolute errors of estimates of *ρ*_*V*_ as the magnitude of the true *ρ*_*V*_ increased. For species with a moderate inter-species correlation (*ρ*_*V*_ *≤* 0.5), and mean probabilities of detection and occupancy of *p* = *m* = 0.5, it was found that *R* = 3 survey replications are required for a reasonable estimate of the correlation (17 percent error). For *ρ*_*V*_ = 0.75, we see that *R* = 5 survey replications are required for estimates with similar error levels. These indicative results may help researchers design surveys for JSDMs.

To understand the relationship between *m, p* and *ρ*_*V*_ above, it is tempting to consider the correlation of the species from the imperfectly detected raw (binary) data, namely *cor*(*Y*_*k*_, *Y*_*l*_), and examine how it relates to the true correlation of occupancy, *cor*(*Z*_*k*_, *Z*_*l*_), see Supplement S5-2. However, both *cor*(*Y*_*k*_, *Y*_*l*_) and *cor*(*Z*_*k*_, *Z*_*l*_) reflect dependencies due to the environmental variables as well as the intrinsic correlation *ρ*_*V*_. For this reason, we had to rely on simulation studies to explore the impact of *N, R, m*_*k*_, and *p*_*k*_ on estimates of *ρ*_*V*_. In another working paper, we have attempted to derive analytical relationships.

The Victorian study of owls and gliders illustrates some of the problems these models face with low site numbers and survey replications. These surveys were originally conducted to study distributions of rare and endangered species, with a sample size of *N* = 202 sites and *R* = 2 or *R* = 3 survey replications, as is typical of many such studies. Our analysis resulted in parameter estimates with wide 95% credible intervals, so only a few covariate parameters were deemed significant. The small number of replications in this dataset, most likely resulted in biased intrinsic correlation estimates due to imperfect detection. To achieve better results, strategies to increase the number of survey replications are required. Previous papers suggest employing multiple independent observers at a site, or conducting multiple surveys during a visit to a site (Mackenzie & Royle 2005) to increase survey replications. Alternatively, researchers could consider time-dependent detection models (Guillera-Arroita *et al*. 2011), though the effectiveness of models of this type with JSDMs needs further investigation. Survey design should also aim to maximise detection probabilities for target species. For instance, surveys using environmental DNA can enhance the detection of some species (Dejean *et al*. 2012).

We also consider if the results were affected by the number of species modelled at one time. Tobler *et al*.(2019) studied multivariate probit JSDMs and their performance over different number of sites for different number of species and concluded that accuracy of parameter estimates decreased for twenty or more species. For our case study, there are only up to four species at any site. Due to the small numbers, there was no significant changes in parameter estimates as the number of species increased from two to four.

The effectiveness of the proposed JSDM explicitly accounting for detection was compared to a model which did not explicitly account for detection but uses collapsed data. We show through simulations that the model using collapsed data is inferior to the JSDM explicitly accounting for detection, when estimating the true probability of occupancy across geophysical or environmental spaces. For collapsed data models, estimates of the probability of joint occupancy (*ω*_11_) can be confounded by covariates of detection and provide an erroneous understanding of species distributions. However, both models give similar results when predicting the observed occupancy. This finding is in line with findings from previous studies of single species distribution models (Lahoz-Monfort *et al*. 2014). These results are applicable to both joint and marginal species distributions. We also show through simulations that estimates of intrinsic correlation between species are similar for the two approaches. This is true even for low values of *N* and *R*, where estimates were similarly poor. This similarity is most likely because the collapsed data has equivalent information to the original survey data with replications as shown by Equation 7.

Our collapsed data results could also apply to JSDMs applied to large scale presence-absence data, where data for grid cells may be drawn from multiple observations over some years (Isaac *et al*. 2020). In such a case, an accumulation of data points collected over time could be considered aggregated in a similar manner to a collapsed dataset. Our work gives an understanding of the impact of imperfect detection on the findings of such models, and is also a warning for the biases that may be introduced.

Imperfect detection has a serious effect on JSDMs, and can result in biased estimates of intrinsic correlation between species and significant errors in predictions of joint (and marginal) probabilities of occupancy. Even models that explicitly account for detection need to be used with care: their results may still be biased, particularly for low probabilities of detection. A key approach to reducing errors and bias is careful study design that aims to select suitable values of *N* and *R* given the target species and environmental conditions of the study area. Information on possible occupancy and detection rates, such as historic data from previous surveys, used in conjunction with simulations such as those in this paper, may prove useful in determining these values. If the probability of detection is low, researchers may find the number of survey replications required is difficult or unrealistic to achieve. In this case, the magnitude of most parameters may be significantly underestimated. However, the problem of bias is not an issue when *N* and *R* are chosen to be sufficiently high.

## Supporting information

Model Code

Further Simulation Results

Further Case Study Results

Sensitivity Analysis

Relations Observed and Actual Data

## Acknowledgements

We gratefully acknowledge the assistance of Dr.Matt White and the Arthur Rylah Institute, Department of Environment, Land, Water and Planning (Victoria) for providing their field data for the Victorian Central Highland Owl and Glider Study. The support of the Australian Research Council grant DPI 150102472 is gratefully acknowledged.

## Authors’ contributions

All analysis was carried out by SH. The underlying theory, design of simulations, and writing of the manuscript was a collaboration between SH, YW, and LS.

## Data accessibility

The presence absence data and covariates used in this analysis are archived and can be found at (dryad archive, address to be determined). R scripts used to generate simulated data can be found in the supplementary information.

## References

Austin, M.P. (2002) Spatial prediction of species distribution: an interface between ecological theory and statistical modelling. Ecological Modelling, 157, 101–118.

Burgman, M.A., Lindenmayer, D.B. & Elith, J. (2005) Managing landscapes for conservation under uncertainty. Ecology, 86, 2007–2017.

Chib, S. & Greenberg, E. (1998) Analysis of multivariate probit models. Biometrika, 85, 347–361.

Comte, L. & Grenouillet, G. (2013) Species distribution modelling and imperfect detection: comparing occupancy versus consensus methods. Diversity and Distributions, 19, 996–1007.

Dejean, T., Valentini, A., Miquel, C., Taberlet, P., Bellemain, E. & Miaud, C. (2012) Improved detection of an alien invasive species through environmental dna barcoding: the example of the american bullfrog lithobates catesbeianus. Journal of Applied Ecology, 49, 953–959.

Demirhan, H. & Hamurkaroglu, C. (2008) Bayesian estimation of log odds ratios from r x c and 2 x 2 x k contingency tables. Statistica Neerlandica, 62, 405–424.

Diaconis, P. & Freedman, D. (1986) On the consistency of bayes estimates. Ann Statist, 14, 1–26.

Dorazio, R.M. (2014) Accounting for imperfect detection and survey bias in statistical analysis of presence-only data. Global Ecology and Biogeography, 23, 1472–1484.

Dorazio, R.M. & Royle, J.A. (2005) Estimating size and composition of biological communities by modeling the occurrence of species. Journal of the American Statistical Association, 100, 389–398.

Dorazio, R.M. & Connor, E.F. (2014) Estimating abundances of interacting species using morphological traits, foraging guilds, and habitat. PLOS ONE, 9, 1–9.

Guillera-Arroita, G., Lahoz-Monfort, J.J., MacKenzie, D.I., Wintle, B.A. & McCarthy, M.A. (2014) Ignoring imperfect detection in biological surveys is dangerous: A response to fitting and interpreting occupancy models’. PLOS ONE, 9, 1–14.

Guillera-Arroita, G., Morgan, B.J.T., Ridout, M.S. & Linkie, M. (2011) Species occupancy modeling for detection data collected along a transect. Journal of Agricultural, Biological, and Environmental Statistics, 16, 301–317.

Guillera-Arroita, G., Ridout, M.S. & Morgan, B.J.T. (2010) Design of occupancy studies with imperfect detection. Methods in Ecology and Evolution, 1, 131–139.

Harris, D.J. (2015) Generating realistic assemblages with a joint species distribution model. Methods in Ecology and Evolution, 6, 465–473.

Hui, F.K.C., Warton, D.I. & Foster, S.D. (2015) Community level models, finite mixture models, penalized likelihood, regularization, species archetype models, variable selection. Annals of Applied Statistics, 9, 866–882.

Hurtado Rúa, S.M., Mazumdar, M. & Strawderman, R.L. (2015) The choice of prior distribution for a covariance matrix in multivariate meta-analysis: a simulation study. Statistics in medicine, 34, 40834104.

Isaac, N.J.B., Jarzyna, M.A., Peter, K., Boersch-Supan, P.H., Browning, E., Freeman, S.N., Golding, N., Guillera-Arroita, G., Henrys, P.A., Jarvis, S., Lahoz-Monfort, J., Pagel, J., Prescott, O.L., Reto, S., Simmonds, E.G. & O’Hara, R.B. (2020) Data integration for large-scale models of species distributionsy. Trends in Ecology and Evolution, 35, 40834104.

Kavanagh, R. (2002) Comparitive Diets of the Powerful Owl (Ninox Strenua), Sooty Owl (Tyto Tenebricosa) and Masked Owl (Tyto Novaehollandiae) in Southeastern Australia, pp. 175–191. CSIRO Publishing.

Kavanagh, R.P. & Bamkin, K.L. (1995) Distribution of nocturnal forest birds and mammals in relation to the logging mosaic in south-eastern new south wales, australia. Biological Conservation, 71, 41–53.

Kéry, M. & Royle, J.A. (2016) Applied Hierarchical Modeling in Ecology. Academic Press, Boston.

Lahoz-Monfort, J.J., Guillera-Arroita, G. & Wintle, B.A. (2014) Imperfect detection impacts the performance of species distribution models. Global Ecology and Biogeography, 23, 504–515.

Lemoine, N.P. (2019) Moving beyond noninformative priors: why and how to choose weakly informative priors in bayesian analyses. Oikos, 128, 912–928.

Lumsden, L.F., Nelson, J.L., Todd, C.R., Scroggie, M.P., McNabb, E.G., Raadik, T.A., Smith, S.J., Acevedo, S., Cheers, G., Jemison, M.L. & Nicol, M.D. (2013) A new strategic approach to biodiversity management. Technical report, Arthur Rylah Institute for Envrionmental Research, Department of Environment and Primary Industries, Heidelberg, Victoria, Australia.

Mackenzie, D.I. & Royle, J.A. (2005) Designing occupancy studies: General advice and allocating survey effort. Journal of Applied Ecology, 42, 1105–1114.

MacKenzie, D.I., Bailey, L.L. & Nichols, J.D. (2004) Investigating species co-occurrence patterns when species are detected imperfectly. Journal of Animal Ecology, 73, 546–555.

MacKenzie, D.I., Nichols, J.D., Lachman, G.B., Droege, S., Royle, J.A. & Langtimm, C.A. (2002) Estimating site occupancy rates when detection probabilities are less than one. Ecology, 83, 2248–2255.

Ovaskainen, O., Abrego, N., Halme, P. & Dunson, D. (2015) Using latent variable models to identify large networks of species-to-species associations at different spatial scales. Methods in Ecology and Evolution, 7, 549–555.

Ovaskainen, O., Hottola, J. & Siitonen, J. (2010) Modeling species co-occurrence by multivariate logistic regression generates new hypotheses on fungal interactions. Ecology, 91, 2514–2521.

Plummer, M. (2016) Bayesian Graphical Models using MCMC. R Foundation for Statistical Computing, Vienna, Austria.

Pollock, L.J., Tingley, R., Morris, W.K., Golding, N., O’Hara, R.B., Parris, K.M., Vesk, P.A. & McCarthy, M.A. (2014) Understanding co-occurrence by modelling species simultaneously with a joint species distribution model (jsdm). Methods in Ecology and Evolution, 5, 397–406.

R Core Team (2015) R: A language and environment for statistical computing. R Foundation for Statistical Computing, Vienna, Austria.

Rota, C.T., Ferreira, M.A.R., Kays, R.W., Forrester, T.D., Kalies, E.L., McShea, W.J., Parsons, A.W. & Millspaugh, J.J. (2016) A multispecies occupancy model for two or more interacting species. Methods in Ecology and Evolution, 7, 11641173.

Royle, J.A. (2004) N-mixture models for estimating population size from spatially replicated counts. Biometrics, 60, 108–115.

Royle, J.A. & Dorazio, R.M. (2008) Hierarchical Modeling and Inference in Ecology: The analysis of data from populations, metapopulations and communities. Academic Press, Elsevier Inc., San Diego, California.

Shoener, T.W. (2009) The ecological niche. The Princeton Guide to Ecology. Princeton University Press.

Thorson, J.T., Scheuerell, M.D., Shelton, A.O., See, K.E., Skaug, H.J. & Kristensen, K. (2015) Spatial factor analysis: a new tool for estimating joint species distributions and correlations in species range. Methods in Ecology and Evolution, 6, 627–637.

Tobler, M.W., Kéry, M., Hui, F.K.C., Guillera-Arroita, G., Knaus, P. & Sattler, T. (2019) Joint species distribution models with species correlations and imperfect detection. Ecology, 100, e02754.

Tong, Y.L. (1990) The Multivariate Normal Distribution. Springer Series in Statistics. Springer New York, New York, NY.

Tyre, A.J., Tenhumberg, B., Field, S.A., Niejalke, D., Parris, K. & Possingham, H.P. (2003) Improving precision and reducing bias in biological surveys: Estimating false-negative error rates. Ecological Applications, 13, 1790–1801.

Waddle, J.H., Dorazio, R.M., Walls, S.C., Rice, K.G., Beauchamp, J., Schuman, M.J. & Mazzotti, F.J. (2010) A new parameterization for estimating co-occurrence of interacting species. Ecological Applications, 20, 1467–1475.

Warton, D. (2015) New opportunities at the interface between ecology and statistics. Methods in Ecology and Evolution, 6, 363–365.

Wintle, B.A., Kavanagh, R.P., McCarthy, M.A. & Burgman, M.A. (2005) Estimating and dealing with detectability in occupancy surveys for forest owls and arboreal marsupials. Journal of Wildlife Management, 69, 905–917.

Wintle, B.A., McCarthy, M.A., Parris, K.M. & Burgman, M.A. (2004) Precision and bias of methods for estimating point survey detection probabilities. Ecological Applications, 14, 703–712.

