## Supplementary material for "Effectiveness of Joint Species Distribution Models in the Presence of Imperfect Detection": Model Code

### Effectiveness of JSDMs Allowing for Detection Supplement 1: Model Extensions and Code

School of Science, RMIT, 124 LaTrobe St, Melbourne, Victoria, 3000, Australia

June 2019

#### 1 Introduction

This supplement provides some minor extensions to the detection and collapsed data models used in the paper *Analysis of JSDMs Subject to Imperfect Detection.* It also provides code for JAGs models used in the paper.

#### 2 Model Extensions

##### 2.1 Modelling Detection

If detection varies with time sensitive covariates, such as wind, time of day or season, then the detection model can be updated as follows:

$$\begin{aligned}
&Y_{ikj} = D_{ikj} Z_{ik} \\
&D_{ikj} \sim Bernoulli(p_{ikj}) \\
&logistic(p_{ikj}) = \alpha_{0k} + W_{ij}^T \alpha_k \\
&W_{ij} = (W_{i1j}, W_{i2j}, .., W_{idj})
\end{aligned} \tag{1}$$

$W_{idj}$ is the vector of d detection covariates for site $i$ observation $j$.

##### 2.2 Collapsed Data Model

If detection covariates vary with time, there may be multiple values of these covariates per site. However, in a collapsed model, only a single value for the detection covariates can be used per site. To incorporate these time varying detection variables in a collapsed model, a summary value is required. For instance, the mean value of continuous a covariate across replications for a site could be used. The collapsed model could then be written as:

$$\mu_{ik}^* = \beta_{0k} + X_i^T \beta_k + W_i^{*T} \beta_k^* \tag{2}$$

where $\mu_{ik}$ is the effective latent occupancy for species k at site i, $\beta_k^*$ is the vector of pseudo slope parameters for the collapsed (or summarized) detection covariates $W_i^*$.

### 3 Implementing the JAGS model

This section provides code for the model explicitly accounting for detection and the model for collapsed data, which does not include a detection component. It then outlines modifications used to look to handle particular cases, including constant detection across sites and constant occupancy across sites.

#### 3.1 Model Explicitly Accounting For Detection

This listing was developed from the community models (Dorazio & Royle 2005; Royle & Dorazio 2008; Kéry & Royle 2016), modifying them to implement the probit occupancy model of Pollock *et al.* (2014). The detection approach is adapted from the model JAGs model proposed by Kéry & Royle (2016) for community models.

```
model {
    # Priors
    # Species-specific effects in occupancy and detection
    for(m in 1:M){  # per species
      # Parameters
      # occupancy
      lpsi[m] ~ dnorm(mu.lpsi, tau.lpsi) # intercept
      for ( i in 1:novar.occ ) {          # covariate parameters
        betalpsi[m,i] ~ dnorm(mu.betalpsi[i], tau.betalpsi[i])
      }
      # detection
      lp[m] ~ dnorm(mu.lp, tau.lp)        # intercept
      for ( i in 1:novar.det ) {
        alphalp[m,i] ~ dnorm(mu.alphalp[i], tau.alphalp[i])
      }
    }

    # Hyperpriors
    # For the model of occupancy
    mu.lpsi ~ dnorm(0,0.01)        # intercept
    tau.lpsi <- pow(sd.lpsi, -2)
    sd.lpsi ~ dunif(0,10) # uniform bounds from trial and error
    for (i in 1:novar.occ) {  # covariate parameters
      mu.betalpsi[i] ~ dnorm(0,0.01)
      tau.betalpsi[i] <- pow(sd.betalpsi[i], -2)
      sd.betalpsi[i] ~ dunif(0, 0.1)
    }
    # correlation component
    Tau[1:M, 1:M] ~ dwish(id[, ], df)
    # For the model of detection
    mu.lp ~ dnorm(0,0.1)         # intercept
    tau.lp <- pow(sd.lp, -2)
    sd.lp ~ dunif(0, 2)
    for ( i in 1:novar.det ) {  # covariate parameters
      mu.alphalp[i] ~ dnorm(0,0.1)
      tau.alphalp[i] <- pow(sd.alphalp[i], -2)
      sd.alphalp[i] ~ dunif(0,10)
    }
```

```
  # Ecological model for true occurrence (process model)
  for (n in 1:nsite) {  # for each site
    V[n, 1:M] ~ dmnorm(mu.occ[n, ], Tau[, ])
    for(m in 1:M){   # for each species
      mu.occ[n,m] <- lpsi[m] + inprod(betalpsi[m,],hab[n,])
      # no occupancy covariates > 1
      z[n,m] ~ dbern(psi[n,m])
      psi[n,m] <- step(V[n,m] )
    }
  }
  # Observation model for replicated
  # detection/nondetection observations
  for (n in 1:nsite){
    for(m in 1:M){  # per species
      for(r in 1:s.reps[n]){
        logit(p[n,r,m]) <-lp[m]+inprod(alphalp[m,],wind[n,r,])
        y[n,r,m] ~ dbern(z[n,m] * p[n,r,m])
      }
    }
  }

  # Derived quantities
  for(m in 1:M){
    Nocc.fs[m] <- sum(z[,m]) # Number of occupied sites
  }
  for (n in 1:nsite){
   Nsite[n] <- sum(z[n,])
   # Number of occurring species at each site
  }
  for ( m in 1:M) {
    for (n in 1:nsite) {
      Nfreq_obs[m,n] <- sum(y[n,1:s.reps[n],m])
      # number obs per species at each site
    }
  }
  index1 <- step(nz-1)
  # step(nz-1) = 1 if nz > 0, 0 o/w - allows nz = 0
}
```

##### 3.2 Model for Collapsed Data without Explicit Detection

This listing is a simplification of the previous listing, it is equivalent to the latent probit occupancy model suggested by Pollock *et al.*2014.

```
model {
  # Priors
  # Species-specific effects in occupancy and detection
  for(m in 1:M){  # per species
    # Parameters
    #occupancy - now observed occupancy (note)
    lpsi[m] ~ dnorm(mu.lpsi, tau.lpsi) # intercept
    for ( i in 1:novar.occ ) {        # covariate parameters
```

```
      betalpsi[m, i] ~ dnorm(mu.betalpsi[i], tau.betalpsi[i])
    }
  }

  # Hyperpriors
  # For the model of occupancy
  mu.lpsi ~ dnorm(0,0.01)        # intercept
  tau.lpsi <- pow(sd.lpsi, -2)
  sd.lpsi ~ dunif(0,10) # uniform bounds from trial and error
  for (i in 1:novar.occ) {  # covariate parameters
    mu.betalpsi[i] ~ dnorm(0,0.01)
    tau.betalpsi[i] <- pow(sd.betalpsi[i], -2)
    sd.betalpsi[i] ~ dunif(0, 10)
  }
  # correlation component
  Tau[1:M, 1:M] ~ dwish(id[, ], df)

  # Ecological model for true occurrence (process model)
  for (n in 1:nsite) {  # for each site
    V[n, 1:M] ~ dmnorm(mu.occ[n, ], Tau[, ])
    for(m in 1:M){    # for each species
      mu.occ[n,m] <- lpsi[m] + inprod(betalpsi[m,], hab[n,] )
      # no occupancy covariatess > 1
      y[n,m] ~ dbern(psi[n,m])
      # observed occupancy - Y data supplied as presumed Z
      psi[n,m] <- step(V[n,m] )
    }
  }

  # Derived quantities
  for(m in 1:M){
    Nocc.fs[m] <- sum(y[,m]) # Number of occupied sites
  }
  for (n in 1:nsite){
   Nsite[n] <- sum(y[n,])
   # Number of occurring species at each site
  }
  index1 <- step(nz-1)
  # step(nz-1) = 1 if nz > 0, 0 o/w - allows nz = 0
}
```

##### 3.3 Modifications of the Code

The above code shows the JAGS models used for multiple occupancy and detection covariates for the model accounting for detection and multiple occupancy covariates for the model ignoring detection. However, a number of studies use the statistically constant where only the intercept is used for both detection and occupancy. Some other studies looked at constant detection with varying occupancy. Further, it was determined that for a single covariate of occupancy, the R to JAGS interface did not implement this is the same way as multiple covariates of occupancy. JAGS does not support conditional statements for control of the program. For this work, a number of working implementations were constructed to support these cases.

**Implementing constant detection:**

- remove *alpha* parameter priors mu.alphalp and tau.alphalp, and their hyper priors
- change the computation of probability of detection to only consider the intercept parameter, where $lp[m] \equiv \alpha_{0m}$

```
logit(p[n,r,m]) <- lp[m] # constant detection
```

**Implementing constant occupancy:**

- remove *beta* parameter priors mu.betalpsi and tau.betalpsi, and their hyper priors
- change the computation of mean latent detection to only consider the intercept parameter, where $lpsi[m] \equiv \beta_{0m}$

```
mu.occ[n,m] <- lpsi[m]
# No occupancy covariates = 0
```

**Implementing occupancy for a single covariate:**

- change the computation of mean latent occupancy to use a habitat (occupancy parameter) vector rather than a matrix

```
mu.occ[n,m] <- lpsi[m] + inprod(betalpsi[m,],hab[n])
# No occupancy covariates = 1
# Jags annoyingly converting mx to vector
```

### References


Dorazio, R.M. & Royle, J.A. (2005) Estimating size and composition of biological communities by modeling the occurrence of species. *Journal of the American Statistical Association*, **100**, 389–398.

Kéry, M. & Royle, J.A. (2016) *Applied Hierarchical Modeling in Ecology.* Academic Press, Boston.

Pollock, L.J., Tingley, R., Morris, W.K., Golding, N., O'Hara, R.B., Parris, K.M., Vesk, P.A. & McCarthy, M.A. (2014) Understanding co-occurrence by modelling species simultaneously with a joint species distribution model (jsdm). *Methods in Ecology and Evolution*, **5**, 397–406.

Royle, J.A. & Dorazio, R.M. (2008) *Hierarchical Modeling and Inference in Ecology: The analysis of data from populations, metapopulations and communities.* Academic Press, Elsevier Inc., San Diego, California.
