## Supplementary material for "Effectiveness of Joint Species Distribution Models in the Presence of Imperfect Detection": Further Simulation Results

### Effectiveness of JSDMs Allowing for Detection Supplement 2: Simulation Studies

School of Science, RMIT, 124 LaTrobe St, Melbourne, Victoria, 3000, Australia

June 2019

This supplement provides further results for studies 1, 2, 3 and 4 for the paper *Analysis of JSDMs Subject to Imperfect Detection.* It also provides the results of a comparison between models accounting for detection, ignoring detection using collapsed data, and ignoring detection using a single observation.

#### 1 Study Design

This paper investigates the performance of the multivariate probit JSDMs accounting for imperfect detection and compares these results with approaches that use collapsed data. We undertook four simulation studies. For the simulation studies, all data was simulated in accordance with the MVP occupancy model (Equation 3, main paper) and Bernoulli detection model (Equation 4, main paper) using R 3.5.1 [1]. Latent occupancy $V$ was simulated by the mvtnorm R Package [2] and the impact of imperfect detection simulated by the R rbinom function [1]. Bayesian analysis was used to estimate models as described in Section 2.5, main paper. We used two species in our simulation studies,but the key results will also apply to multiple species. For convenience, the covariance matrices used for studies are equivalent to the correlation matrices, with variance of 1 and a single covariance equal to $\rho_V$. Multiple independent data sets were simulated for all parameter combinations, similar to MacKenzie et al. (2002).

**Scenario 1: Determine impact of number of sites and replications on parameter estimates.** For this study, data was simulated using the occupancy model (Eq 3, main paper) and detection model (Eq 5, main paper) for varying combinations of site numbers ($N$), survey replications ($R$), and different covariate relations for each species. All covariate data was generated using a uniform distribution between -1 and 1. Data sets, including covariates, were generated independently for 5 simulations for each combination of $\rho_V$, $N$ and $R$ , where $\rho_V \in \{-0.5, 0, 0.5, 0.9\}$, $N \in \{50, 100, 200, 400\}$ and $R \in \{3, 5, 7\}$. Model fitting assumed the same occupancy and detection covariates as the simulated data. For results for varying N and R shown in Section 1.2 of this supplement, we used a single occupancy covariate ($X_1$) and one detection covariate ($W_1$), which are independent. The slope parameters were set to unity for species 1 ($\beta_{11} = 1$, $\alpha_{11} = 1$), and to the negative one for species 2 ($\beta_{12} = -1$, $\alpha_{12} = -1$).

As well, we compare a number of cases for $N = 200$ and $R = 3$ for a number of different covariate cases that cover both dependent occupancy and detection covariates and independent occupancy and detection covariates. Dependent cases arise where $X_1$ and $W_1$ covariates are correlated with each other. For example, if species occupancy depended on vegetation of a certain type while density of vegetation affected detection of the species.

Dependent variables are said to be positively correlated if the detection and occupancy for a species vary with the covariate in the same way, i.e. both positively (increasing as the covariate increases) or both negatively (deceasing as the covariate decreases). Dependent variables are said to be negatively correlated when detection and occupancy for a species vary in opposite ways (i.e. one positively and the other negatively).

The independent cases show how model parameter estimates are impacted when occupancy and detection vary positively and negatively with independent covariates. The following cases are shown:
Case 4: occupancy and detection vary positively, $\beta_{11} = 1$, $\alpha_{11} = 1$;
Case 5: occupancy varies negatively while detection varies positively, $\beta_{11} = -1$, $\alpha_{11} = 1$;
Case 7: occupancy varies positively while detection varies negatively, $\beta_{11} = 1$, $\alpha_{11} = -1$;
Case 8: occupancy and detection vary negatively, $\beta_{11} = -1$, $\alpha_{11} = -1$.

The dependent cases show how model parameter estimates are impacted when occupancy and detection vary positively and negatively with covariates, as follows:
Case 13: occupancy and detection vary positively (positively correlated), $\beta_{11} = 1$, $\alpha_{11} = 1$;
Case 14: occupancy varies negatively while detection varies positively (negatively correlated), $\beta_{11} = -1$, $\alpha_{11} = 1$;
Case 15: occupancy varies positively while detection varies negatively (negatively correlated), $\beta_{11} = 1$, $\alpha_{11} = -1$;
Case 16: occupancy and detection vary negatively (positively correlated), $\beta_{11} = -1$, $\alpha_{11} = -1$.
Case 13 and 16 show positive correlation between the occupancy and detection variables (i.e. both vary in the same direction) while case 14 and 15 show negative correlation between the occupancy and detection variables (i.e. they vary in opposite directions). For these simulations, dependency is shown by setting $X_1 = W_1$, so the occupancy and detection variables coincide.

**Scenario 2: Determine the impact of occurrence and detection probability on $\rho_V$ estimates.** For this study, data was simulated assuming species had identical mean probabilities of detection $p = p_1 = p_2$ and probabilities of occurrence $m = m_1 = m_2$. Occupancy data was simulated using Equation 3 where $m = \phi(\mu) = \phi(\beta_0)$, and detection simulated using Equation 5 where $p = logit^{-1}(\alpha_0)$. Data sets were generated independently for 14 simulations over $N = 200$ sites of each combination of $p$,$m$, $\rho_V$, and $R$. We chose the following values: $p \in \{0.25, 0.5, 0.75, 0.9\}$, $m \in \{0.2, 0.5, 0.8, 0.85\}$ ($\mu \in \{-0.83, 0, 0.83, 1.04\}$), $\rho_V \in \{0, 0.25, 0.5, 0.75, 0.9\}$. $\rho_V$ was estimated using Bayesian regression analysis for $R \in \{3, 5, 7, 10\}$. As well, $\rho_V$ was estimated for a single survey replication, with no detection component (i.e. a JSDM with just the occupancy component).

The study was also extended to include negative intrinsic correlation values, for the following values: $\rho_V \in \{-0.25, -0.5, -0.75\}$, $p \in \{0.25, 0.5, 0.75\}$, and $m = 0.5$, where both species had the same occupancy and detection properties.

**Scenario 3: Compare effectiveness for JSDM explicitly accounting for detection for estimating parameters compared to collapsed data approach.** Using the simulation data from the previous study, collapsed observation data was generated for each simulation. $\rho_V$ and other parameters were then estimated using the collapsed model and Bayesian analysis. Parameter estimates for each $N$ and $R$ and covariate combination were then compared between the model explicitly accounting for detection and the model using collapsed data.

**Study 4: Determine the effectiveness of explicit detection in a JSDM for**

**estimates of joint probability.** $\omega_{11}$ represents the probability of two species occurring together at a site. This measure varies spatially and is affected by both intrinsic correlation between species and environmental conditions. We calculated true values of $\omega_{11}$ using the true $\rho_V$ and occupancy parameters as $W_1$ and $X_1$ varies from -1 to 1. This is then compared to estimated values of $\omega_{11}$ calculated from mean estimated values of $\rho_V$ and other parameters for both the model explicitly accounting for detection (study 2) and the model using collapsed data (study 3). True and predicted values of $\omega_{11}$ values are plotted using colour coded 2D plots.

We also compared estimates of $\omega_{11}$ for a JSDM with detection against approximations of $\omega_{11}$ for single species models, which assume there is no intrinsic correlation between species. So far we have only considered JSDMs which account for dependencies between species. These models provide insights into intrinsic correlation and joint species distributions. Comparing joint probabilities calculated by JSDMs to approximations of single species models gives insight into the impact of intrinsic correlation on species distributions. To do this, we looked at two single species distribution models, one based on a commonly used logit approach [4] and the other using a probit approach. Both models incorporate detection.

#### 1.1 Single Species Models for Scenario 4

Using our previous notation, the Logit based single species occupancy model with detection can be written as:

$$\begin{aligned} &Y_{ijk} = D_{ijk} Z_{ik} \\ &D_{ijk} \sim Bernoulli(p_{ik}) \\ &logit(p_{ik}) = \alpha_{0k} + W_i^T.\alpha_k \\ &Z_{ik} \sim Bernoulli(m_{ik}) \\ &logit(m_{ik}) = \beta_{0k} + W_i^T.beta_k \end{aligned} \tag{1}$$

The probit based single species occupancy model with detection can be written as:

$$\begin{aligned} &Y_{ijk} = D_{ijk} Z_{ik} \\ &D_{ijk} \sim Bernoulli(p_{ik}) \\ &logit(p_{ik}) = \alpha_{0k} + W_i^T.\alpha_k \\ &P(Z_{ik} \geq 0) = P(V_{ik} = 1) \\ &V_{ik} \sim N(X_i^T \beta_k, \sigma_k) \\ &m_{ik} = \beta_{0k} + W_i^T.beta_k \end{aligned} \tag{2}$$

Both models were used to analyse a subset of the simulated data generated for study 2. estimates of other parameters were made with the model using collapsed data for each simulation. These estimates were then compared to those using the explicit model (study 2). Results from these models are compared to the JSDM explicitly accounting for detection, see *"Effectiveness of Joint Species Distribution Models in the Presence of Imperfect Detection"*, equation 8.

#### 1.2 Scenario 1,3, and 4 Simulation Details

Data was simulated using the occupancy model with a single covariate $X_1$ and detection model with a single covariate $W_1$. All covariate data was generated using a uniform distribution between -1 and 1 [1]. Covariates were generated independently for each simulation.

The following occupancy cases were considered:

- Case I: $\beta_{11} = \beta_{21} = 1$ - mean occupancy for both species $\mu = \mu_k$ increases with $X_1$;
- Case II: $\beta_{11} = 1$, $\beta_{21} = -1$ - $\mu_1$ increases with $X_1$ while $\mu_2$ decreases.

These were combined with the following detection cases:

- Case i: $\alpha_{10} = \alpha_{20} = 0$ - constant probability of detection $p = p_k = 0.5$ for both species.
- Case ii: $\alpha_{11} = \alpha_{21} = 1$ - probability of detection for both species $p_k$ increases with $W_1$;
- Case iii: $\alpha_{11} = 1$, $\alpha_{21} = -1$ - $p_1$ increases with $W_1$ while $p_2$ decreases.
- Case iv and v: the same parameters as case 1 and 2, but with $W_1 = X_1$ - i.e. occupancy and detection are dependent.

For all cases the intercept parameters used were $\beta_{01} = \beta_{02} = 0$ for occupancy (equivalent to mean probability of occupancy of 0.5 for each species); and $\alpha_{01} = \alpha_{02} = 0$ for detection (equivalent to mean probability of detection of 0.5 for each species)

#### 2 Scenario 1 results

Estimates of $\rho_V$ and other model parameters are similar between the covariate scenario shown here ($\beta_{11} = \alpha_{11} = 1$ for species 1 and $\beta_{12} = \alpha_{12} = -1$ for species 2) and covariate scenario shown in the main paper ($\beta_{1k} = \alpha_{1k} = 1$ for both species).

Figure 1 shows estimates of $\rho_V$ for this alternate scenario, with R increasing from left to right and N increasing from top to bottom. Each plot shows the estimates for 5 independent simulation for four $\rho_V$ cases, with case values shown by the X axis. As before, we see that bias, the distance of mean values (red circles) from true values (blue triangles), decreases as R increases. We see the uncertainty, relating to the critical interval and range of mean estimates, decreases as N increases. Increasing R may also cause minor decreases in uncertainty.

Figure 2 shows results of estimates for other model parameters. The top two rows show estimates for $\beta_{1k}$, while the bottom two rows show estimates for $\alpha_{1k}$. Note for this alternate study, $\beta_{1k}$ and $\alpha_{1k}$ vary between species: for species one $\beta_{11} = \alpha_{11} = 1$; while for species two $\beta_{21} = \alpha_{21} = -1$. Estimates still work in a similar way to the first scenario, as long as we consider estimates in terms of magnitudes. So the magnitude of $\beta_{1k}$ slope parameters tend to be under estimated, and this bias decreases with R. Similarly, the magnitude of $\alpha_{1k}$ slope parameters tend to be under estimated, and this bias decreases with R. For $\beta_{k0}$ intercept parameters the value is underestimated while for $\alpha_{k0}$ intercept parameters the value is overestimated. Uncertainty in parameter estimates decreases as N increases, and also decrease slightly as R increases.

Estimates of $\rho_V$ and other model parameters are similar between the covariate scenario shown here ($\beta_{11} = \alpha_{11} = 1$ for species 1 and $\beta_{12} = \alpha_{12} = -1$ for species 2) and covariate scenario shown in the main paper ($\beta_{1k} = \alpha_{1k} = 1$ for both species).

Figure 3 compares parameter estimates over number of independent cases (covariate cases 4, 5, 7 and 8; top and third rows) and dependent cases (covariate cases 13, 14, 15 and 16; second row and bottom rows) for $R = 3$ and $N = 200$. Estimates of intrinsic correlation $\rho_V$ show little variation between the different cases as shown by the top and second rows. However, estimates of the $\beta$ occupancy parameters and $\alpha$ detection parameters show some differences. For the independent covariate cases, as shown by third row,

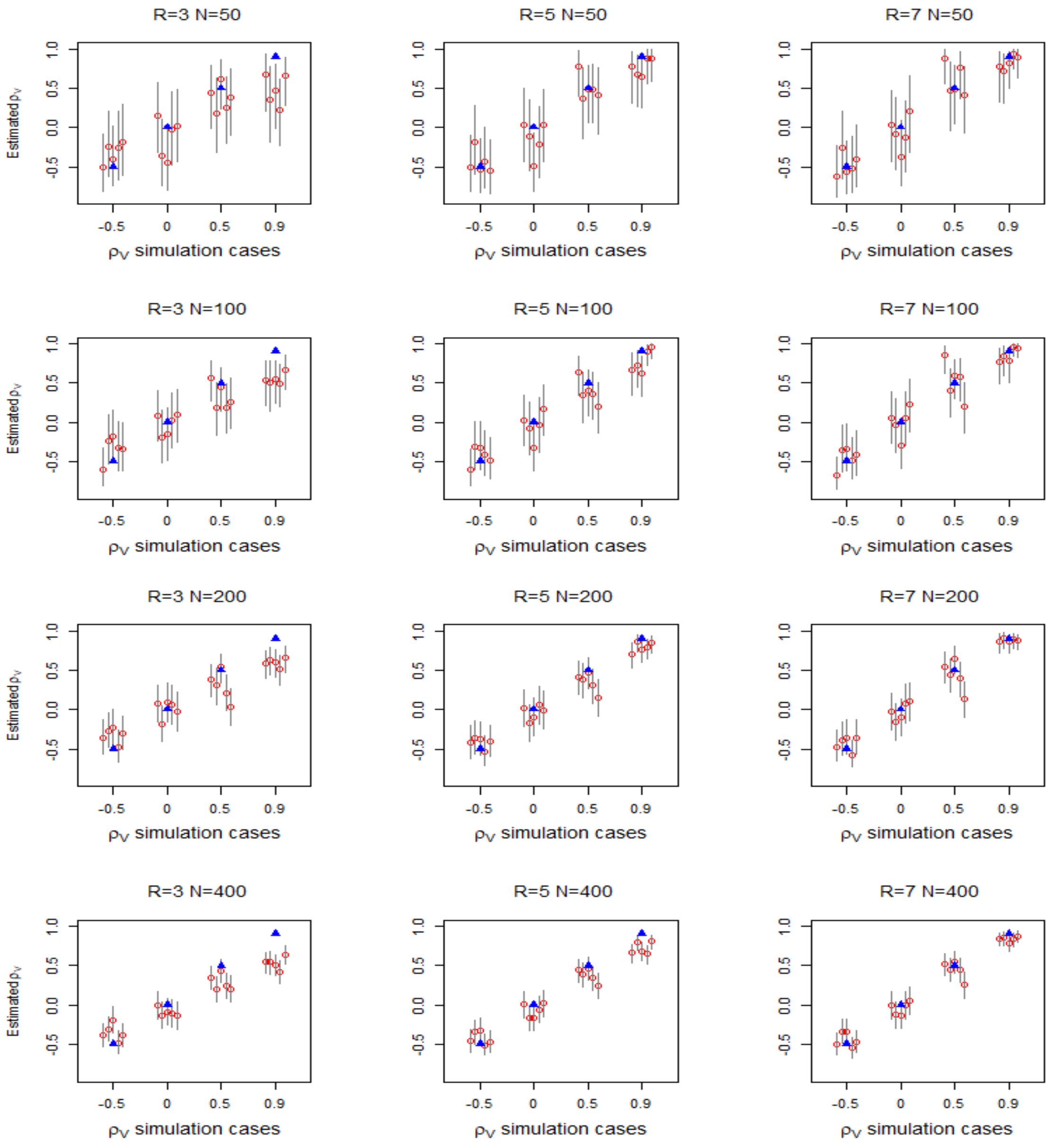


Figure 1: *Estimates of $\rho_V$ for different combinations of number of sites $N$ and number of replications $R$ for species with different occupancy and detection covariate relations. Each panel shows outcomes for one combination of $N$ and $R$ for a number of $\rho_V$ cases, $\rho_V \in \{-0.5, 0, 0.5, 0.9\}$. For each $\rho_V$ case, we show results for 5 simulations as follows: true $\rho_V$ (blue triangle), mean estimate (red circle), and 95% credible intervals limits (grey lines). $N$ increases by row, from top (50) to bottom (400). $R$ increases by column from left (3) to right (7).*

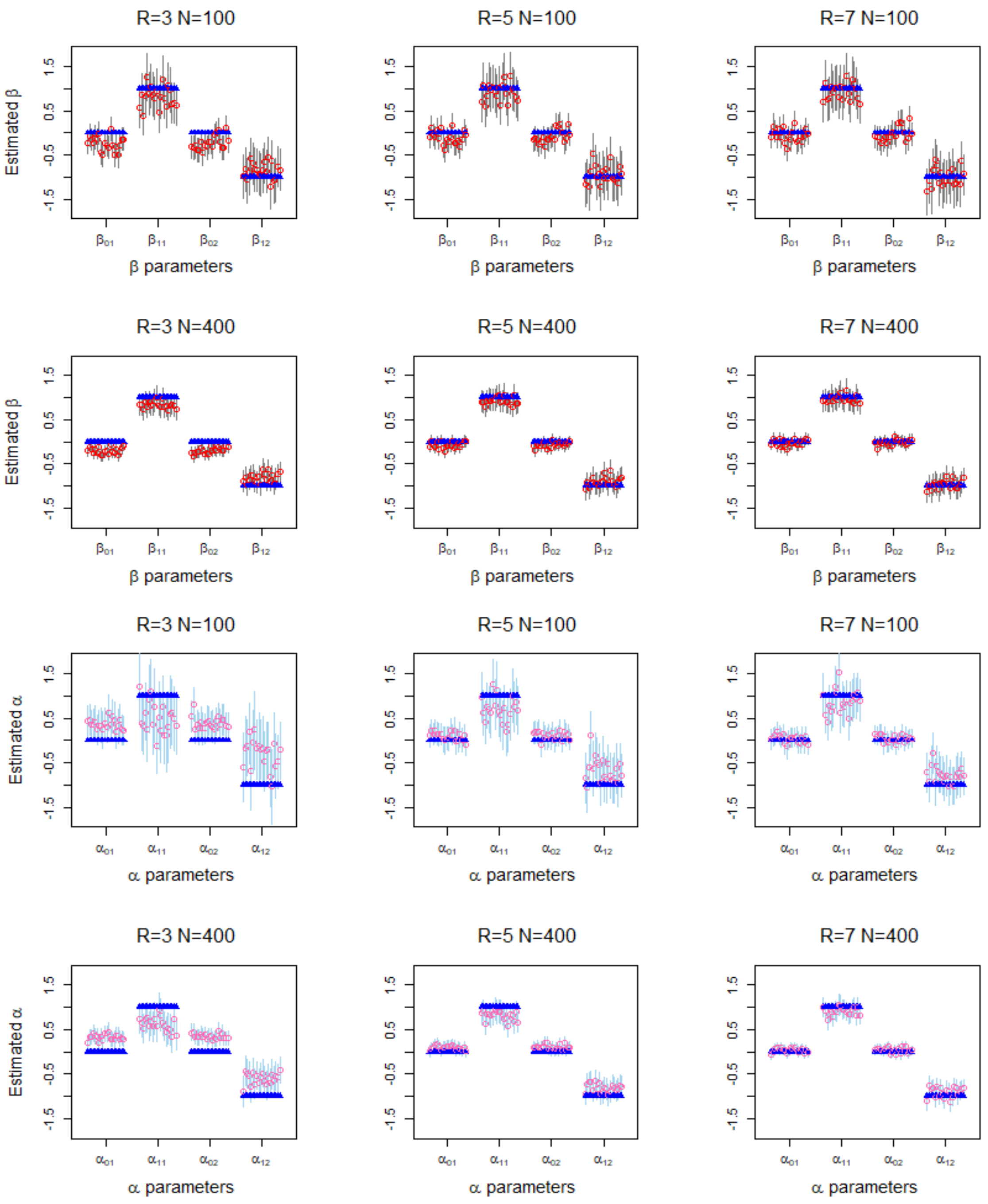


Figure 2: *Impact of survey replications $R$ on estimates of occupancy and detection parameters for species with different occupancy and detection covariate relations. Panels show parameter estimates for $R$ increasing from left (3) to right (7) for* $N = 400$. *Each panel shows 20 estimates for each $\beta$ and $\alpha$ parameter as follows: true value (blue triangle), mean estimate (red circle), and 95% credible intervals limits (grey lines) for model accounting for detection.*

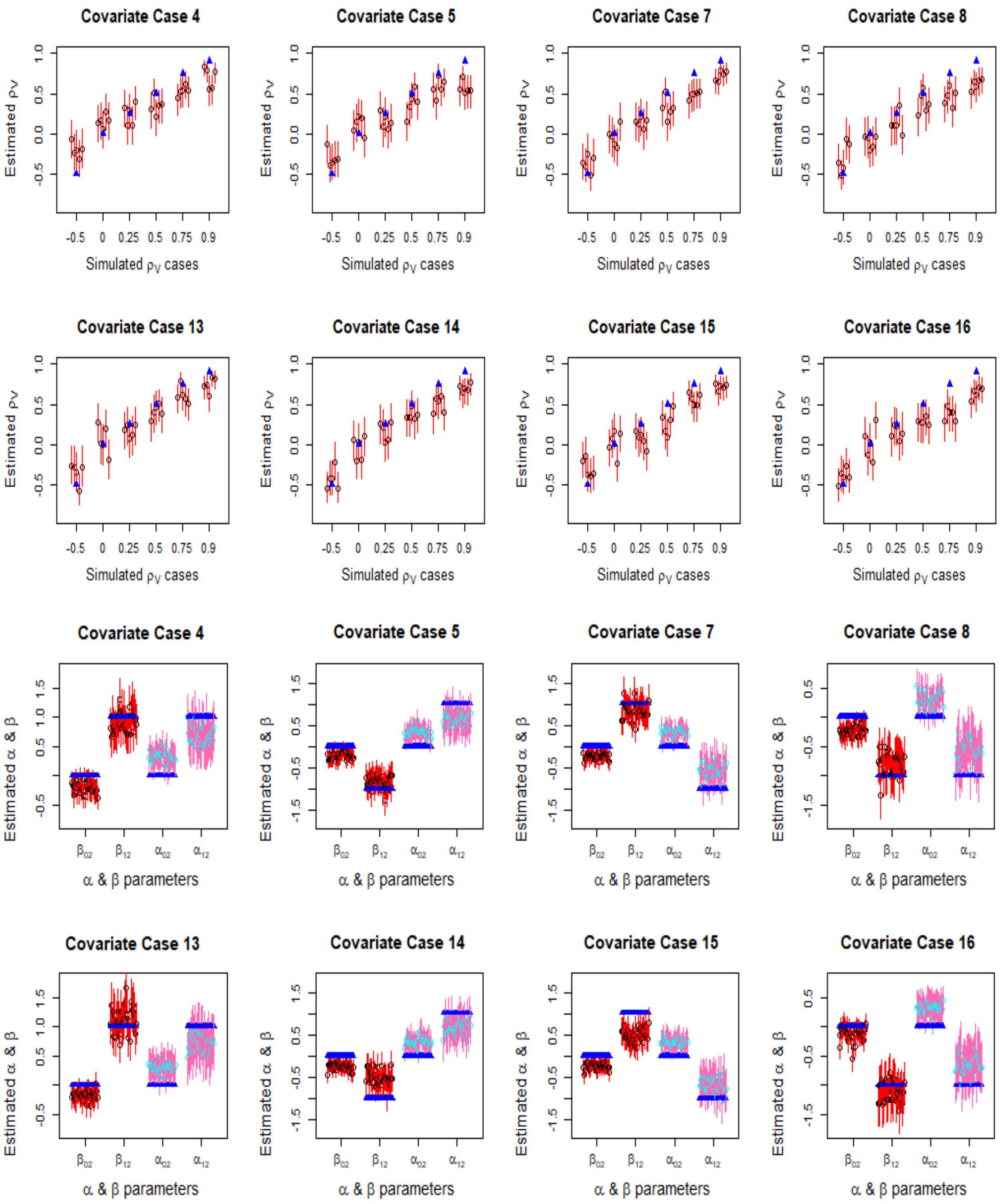


Figure 3: *Comparing model parameters estimates for independent covariate cases (4, 5, 7 and 8) versus dependent covariate cases (13, 14, 15, and 16). Estimates for independent cases are shown by the top and third row; while estiamtes for teh dependent cases are shown for the second and bottom rows. Panels show parameter estimates for each covariate and parameter case for number of sites* $N = 200$ *and replications* $R = 3$*. Each panel shows 20 estimates for each* $\beta$ *and* $\alpha$ *parameter as follows: true value (blue triangle), mean estimate (red circle), and 95% credible intervals limits (grey lines) for model accounting for detection.*

the magnitude of all occupancy parameter estimates is under estimated, as is the magnitude of the detection slope parameter estimate. The bottom rows shows that dependent covariates are positively correlated (covariate cases 13 and 16), the magnitude of occupancy slope parameters tend to be over estimated, as shown by estimates of $\beta_{12}$. The magnitude of occupancy intercept and detection slope parameter however remains under-estimated. Where dependent covariates are negatively correlated (covariate cases 14 and 15), however maintain a similar patter to the independent case. The bias of occupancy slope parameters estimates is slightly larger for the negatively correlated dependent case than the independent case.

### 3 Scenario 2 results

**Scenario 2: The impact of occurrence and detection probability on $\rho_V$ estimates.**

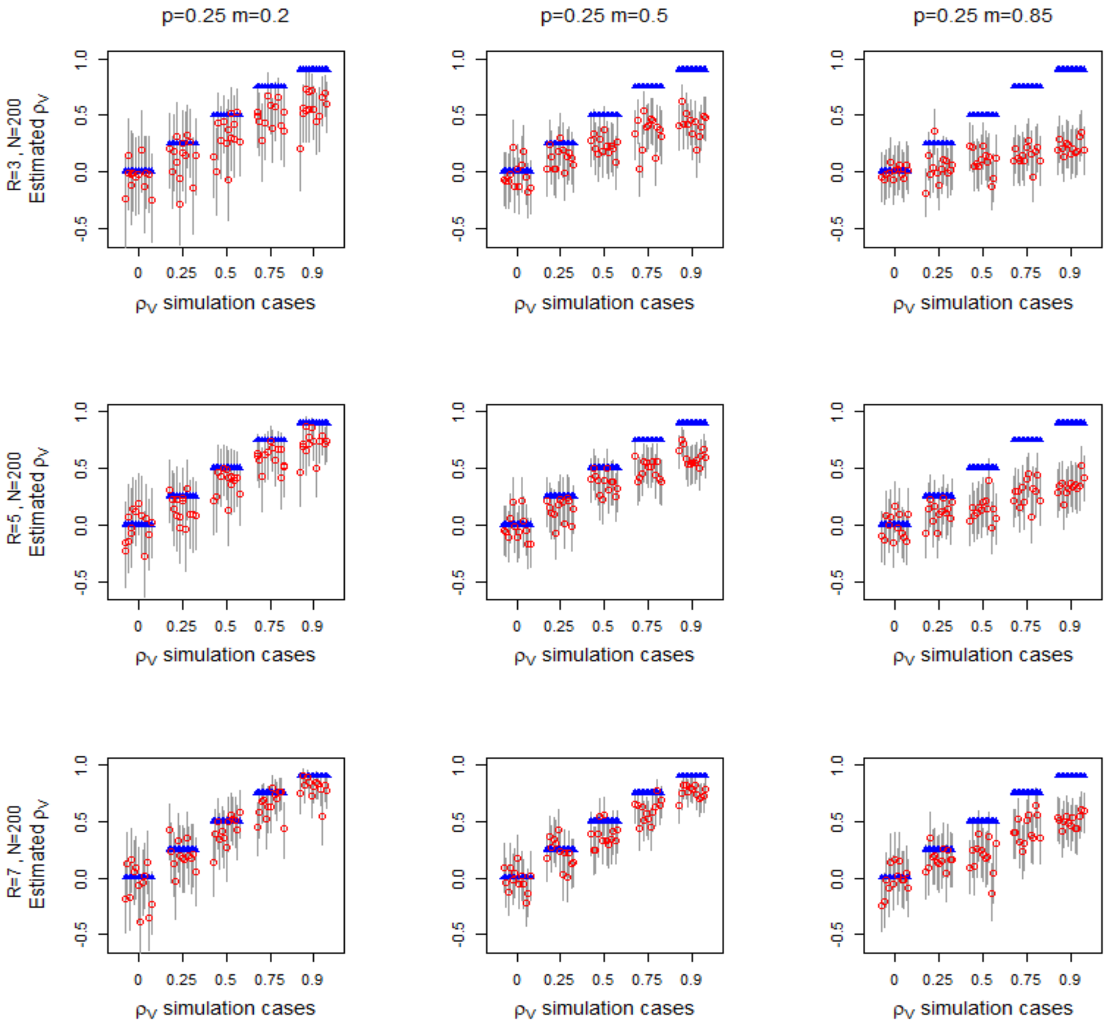


Figure 4: *Estimates of $\rho_V$ for different combinations of probability of detection (p), marginal detection (m), and survey replications (R) where both species have identical properties. 14 independent data sets generated for each combination of p, m, R and $\rho_V$ for 200 sites. Each panel shows 14 results for each $\rho_V$, $\rho_V \in \{0, 0.25, 0.5, 0.75, 0.9\}$, as follows: true $\rho_V$ (blue triangle), mean estimate (red circle), and 95% credible intervals limits (grey lines). Each column shows results for one R value, while each row shows results for one p and m combination.*

Figure 4 rows shows estimates of intrinsic correlation $\rho_V$ increase with $p$ for $m = 0.5$.

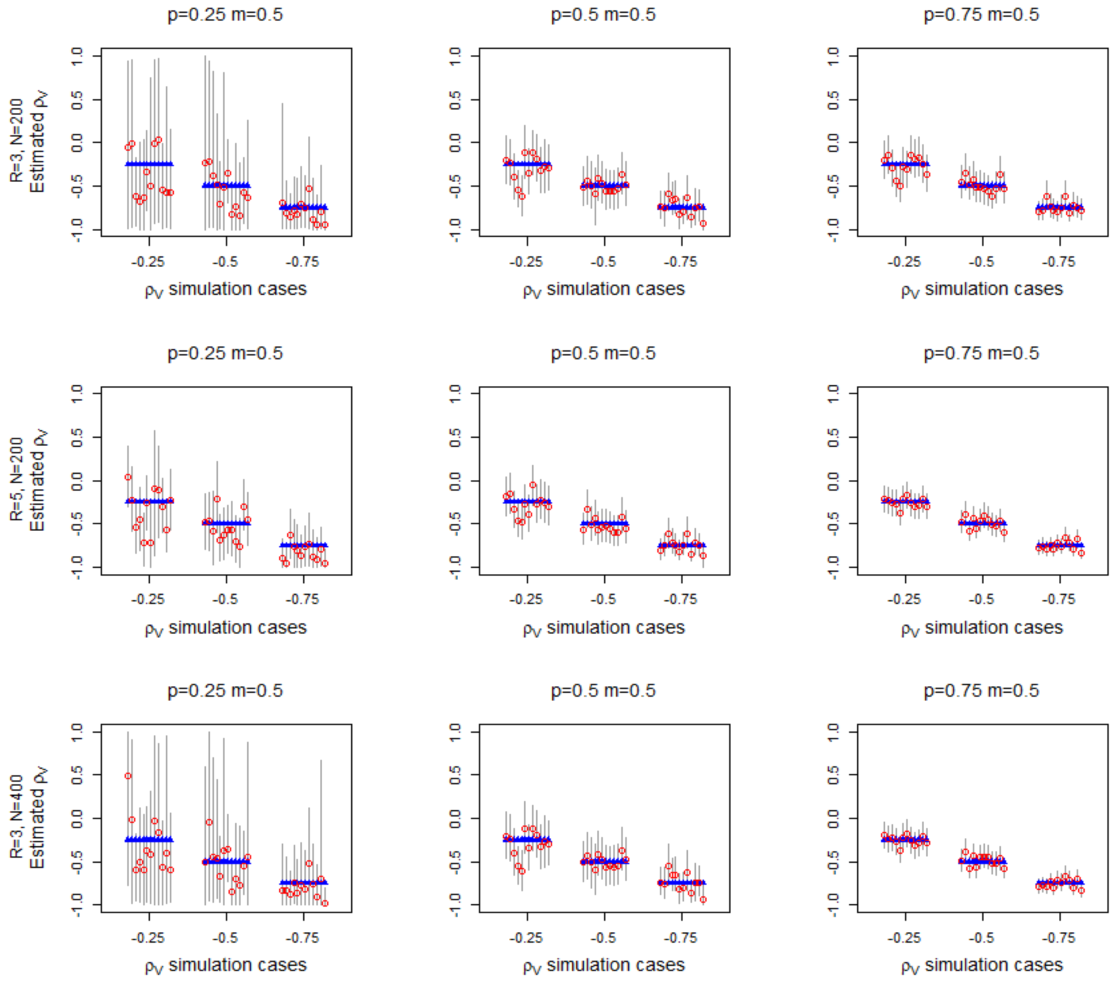


Figure 5: *Estimates of negative intrinsic correlation between species $\rho_V$ for the model explicitly accounting for detection. Estimates are shown for different values of detection probability (p) for marginal occupancy probability $m = 0.5$ where both species have identical properties. Each panel a different p value. 14 independent data sets generated for each combination of p and $\rho_V$ for 200 sites and 3 replications. Each panel shows 14 results for each $\rho_V$, $\rho_V \in \{0, 0.25, 0.5, 0.75, 0.9\}$, as follows: true $\rho_V$ (blue triangle), mean estimate (red circle), and 95% credible intervals limits (grey lines).*

Each plot shows results for 14 simulations for each $\rho_V$ case, R replications and 200 sites. Mean estimates of $\rho_V$ (red circles) are better if they are closer to true $\rho_V$ (blue triangles). Estimates of $\rho_V$ improve as $p$ increases across each row.

*As survey replications $R$ increases fit improves, but high $R$ may be needed for good fit.* Figure 4 shows estimates of $\rho_V$ for $R = 3$ (top row), $R = 5$ (middle row) and $R = 7$ (bottom row). Estimates improve from top to middle row and middle to bottom row, with mean estimates closer to the true values (blue triangles). Increasing $R$ can compensate for lower value of $p$, so estimates of $\rho_V$ for $p = 0.5$ and $R = 5$ are close to estimates of $\rho_V$ for $p = 0.75$ and $R = 3$. Estimates for $p = 0.25$ also improve as $R$ increases from 3 to 5, but further repetitions are required before they match the $p = 0.5$ estimates. Uncertainty also decrease to some extent with increasing R, but remains an issue for low $p$ and $m$ combinations.

*Bias errors are more evident as $\rho_V$ increases.* Bias is generally not apparent for $\rho_V = 0$, but these estimates may be still affected by uncertainty. Uncertainty tends to decrease for

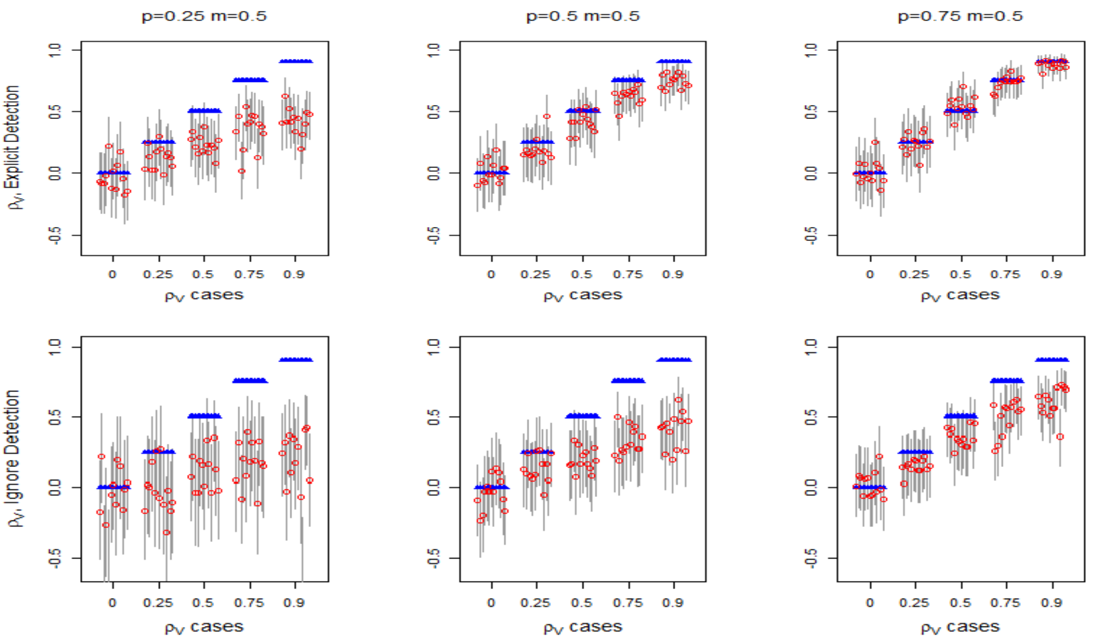


Figure 6: *Estimates of $\rho_V$ for the model explicitly accounting for detection (top row) versus a model completely ignoring detection. Models are compared for different combinations of probability of detection (p), marginal detection (m) where both species have identical properties. Each column shows one p and m combination. 14 independent data sets generated for each combination of p, m, and $\rho_V$ for 200 sites and 3 replications. Each panel shows 14 results for each $\rho_V$, $\rho_V \in \{0, 0.25, 0.5, 0.75, 0.9\}$, as follows: true $\rho_V$ (blue triangle), mean estimate (red circle), and 95% credible intervals limits (grey lines).*

high values of $\rho_V$.

Figure 6 compares estimates of $\rho_V$ for a model explicitly allowing for detection to those from a model completely ignoring detection. These latter models do not include a detection component and use data from a single survey without replications. Each row shows estimates for 14 simulations for each $\rho_V$ case increasing $p$ (left to right) for $m = 0.5$. The top row shows the results for the model allowing for detection with 3 replications and 200 sites, while the bottom row shows the model ignoring detection.

*For negative $\rho_V$, overall errors decrease as probability of detection increases.* Figure 5 shows estimates of intrinsic correlation for $N = 200$ and $R = 3$ (top row), $N = 200$ and $R = 5$ (middle row), and $N = 400$ and $R = 3$ (bottom row). We see that for low probability of detection $p = 0.25$: precision is very poor, estimates are generally inaccurate, and there is a tendency to underestimate correlation. Estimates of $\rho_V$ improve as the probability of detection increases and for high probability of detection $p = 0.75$, precision is good and estimates of $\rho_V$ are close to the true value. Increasing survey replication improves estimates: for $R = 5$ uncertainty decreases markedly for low probability of detection and there is some improvement in mean estimate as shown by the middle row. However increasing the number of sites to $N = 400$ made little improvement on uncertainty or accuracy of estimates compare to $N = 200$.

*Model explicitly allowing for detection is more accurate than the model ignoring detection.* Even for a low number of replications (3), the model explicitly allowing for detection is both less biased, i.e. means estimates generally closer to the true value, and shows less uncertainty, i.e. narrower critical intervals. The model explicitly allowing for detection is also more likely to find significant intrinsic correlations $\rho_V$ than the model ignoring detection.

### 4 Scenario 3 results

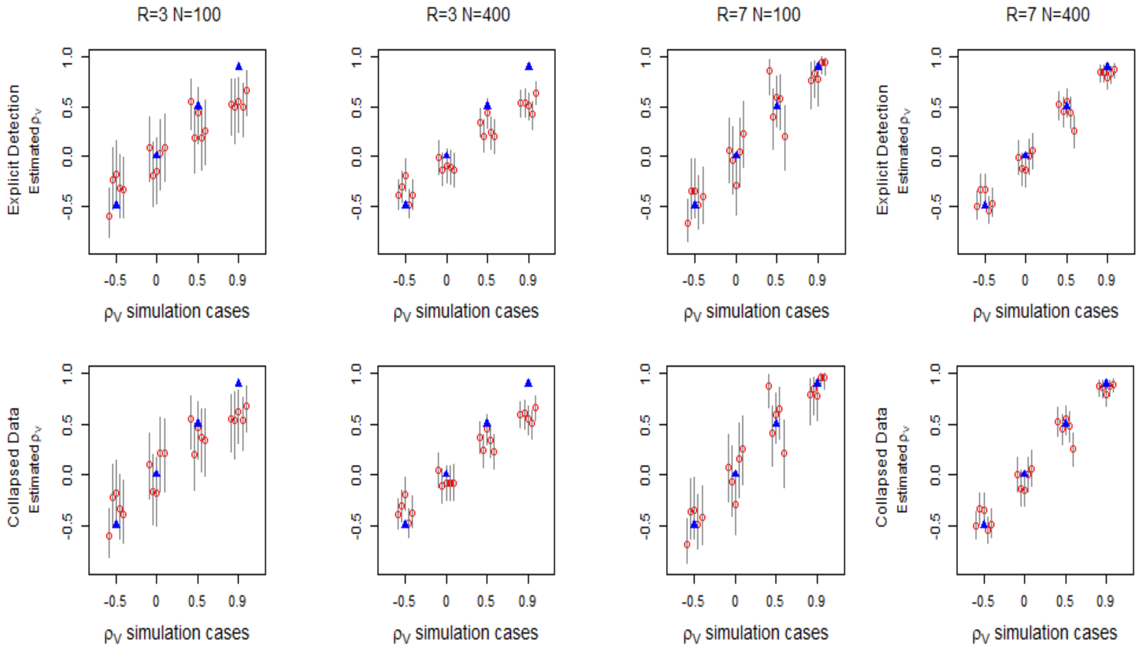


Figure 7: *Impact of using collapsed data to estimate $\rho_V$ for varying number of sites $N$ and survey replications $R$. Top row shows estimates for the model accounting for detection. Bottom row shows estimates for the model using collapsed data for equivalent $N$ and $R$. Each panel shows outcomes for one combination of $N$ and $R$ for a number of $\rho_V$ cases, $\rho_V \in \{-0.5, 0, 0.5, 0.9\}$. For each $\rho_V$ case, we show results for 5 simulations as follows: true $\rho_V$ (blue triangle), effective $\hat{\rho_V}$ (cyan cross), mean estimate (red circle), and 95% credible intervals limits (grey lines).*

Using the collapsed model to estimates of $\rho_V$ and other model parameters has a similar outcome for the covariate scenario shown here ($\beta_{11} = \alpha_{11} = 1$ for species 1 and $\beta_{12} = \alpha_{12} = -1$ for species 2) and covariate scenario shown in the main paper ($\beta_{1k} = \alpha_{1k} = 1$ for both species).

Figure 7 shows estimates of $\rho_V$ for the alternative covariate scenario ($\beta_{11} = \alpha_{11} = 1$, $\beta_{21} = \alpha_{21} = -1$) for both model accounting for detection (top row) and model using collapsed data (bottom row). As with the covariate scenario in the main paper, its shows that estimates of $\rho_V$ remain very similar for both models. Both models show bias decreases as $R$ increases and that uncertainty (width of the credible interval) decreases as $N$ increases. Estimates for the same combination of $N$ and $R$ show little difference between the models.

Figure 7 shows estimates of $\beta_1 k$ and $\alpha_1 k$ parameters for the alternative covariate scenario for both model accounting for detection (top row) and model using collapsed data (bottom row). For the detection model, estimated magnitudes of the parameters tend to be less than the magnitude of the true values for low $R$. Parameter estimates converge to their true values as $R$ increases. A similar result emerges for $\beta_{1k}$ parameters for both species. For $\beta_{2k}$ parameters, which relate to the detection covariates included in the occupancy model, we see that estimates generally become closer to 0 as $R$ increases. Estimates of these parameters do tend have a smaller magnitude than true occupancy covariates. If the original detection relation was negative (e.g. $\alpha_{1k} = -1$) estimates increase as $R$ increases, but if the original detection relation was negative (e.g. $\alpha_{1k} = 1$) estimates decrease as $R$ increases. Uncertainty for low value of $N$ also affects these estimates, $\beta_{2k}$ parameters in particular may not be significant for low values of $N$.

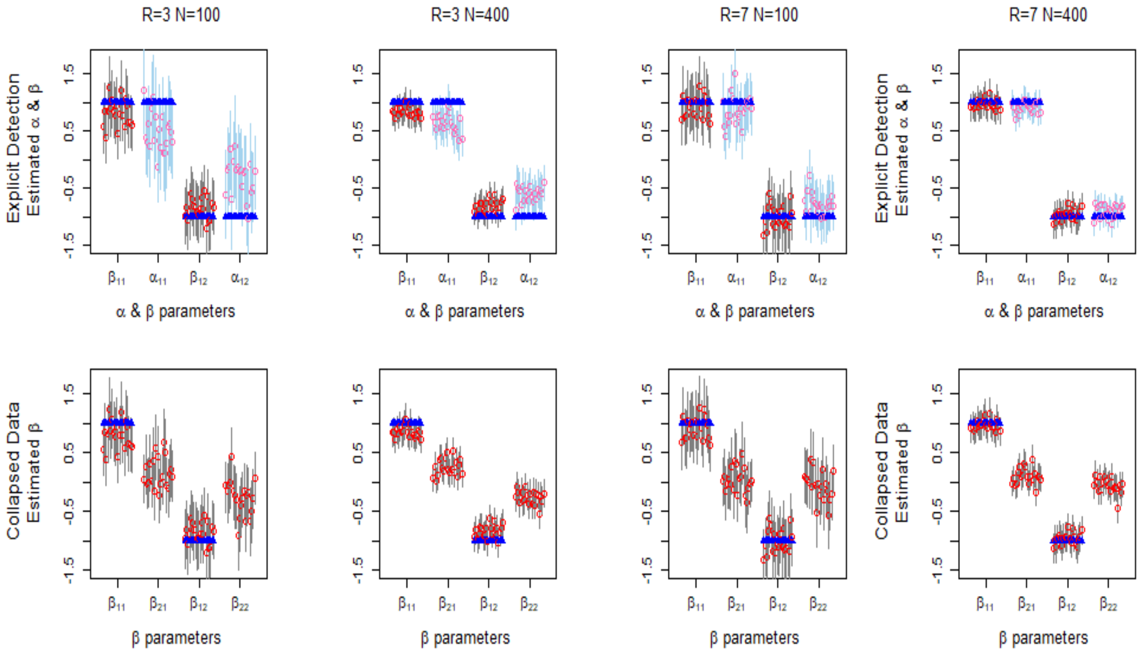


Figure 8: *Impact of using collapsed data to estimate occupancy parameters for varying number of sites N and survey replications R, when species have differing relation to covariates. Top row shows estimates for the model accounting for detection. Bottom row shows estimates for the model using collapsed data for equivalent N and R. Each panel shows 20 estimates of each $\beta$ or $\alpha$ parameter as follows: true value (blue triangle), mean estimate (red/pink circles), and 95% credible intervals limits (grey/blue lines). For* $\beta_{2k}, truevaluesetto0$.

### 5 Scenario 4 results: Comparison of JSDM with explicit detection with JSDM using collapsed data.

Figure 9 shows $\omega_{11}$ for changing $X_1$ and $W_1$ for the alternative covariate scenario ($\beta_{11} = \alpha_{11} = 1$, $\beta_{21} = \alpha_{21} = -1$). Here, $\omega_{11}$ is colour coded, see key on right hand side of each row. The first column shows true $\omega_{11}$, middle column shows estimated $\omega_{11}$ for the model using detection, and the right column shows estimated $\omega_{11}$ for the model using collapsed data. The left column shows that $\omega_{11}$ increase with $X_1$ until $X_1 = 0$ and declines as $X_1$ increases further. Rows show how estimates $\omega_{11}$ for the two models change with N (number of sites) and R (number of replicates).

Estimates of $\omega_{11}$ for the model accounting for detection tend to improve as both N and R increase, with models for low N tending to show positive bias (i.e. not centered round $X_1 = 0$. As N and R increase, estimates become centered closer to $X_1 = 0$, but we note that there are discrepancies even for N=400 and R=7 due to uncertainty.

Estimates of $\omega_{11}$ for the model using collapsed data show the influence of $W_1$. As for the covariate scenario, $W_1$ confounds the estimates, though here the patterns are more variable. Estimates show some improvement as N and R both increase, with higher N reducing bias (offset from 0). Errors are still apparent, due to uncertainty in parameter estimates.

### 6 Scenario 4 results: JSDM and Single Species comparisons

Figure 10 shows estimates of the $\beta$ occupancy parameters and $\alpha$ detection parameters for a single species for three models, the multivariate JSDM (top), single species logit (middle) and single species probit (bottom). The multivariate JSDM estimates are more sensitive to

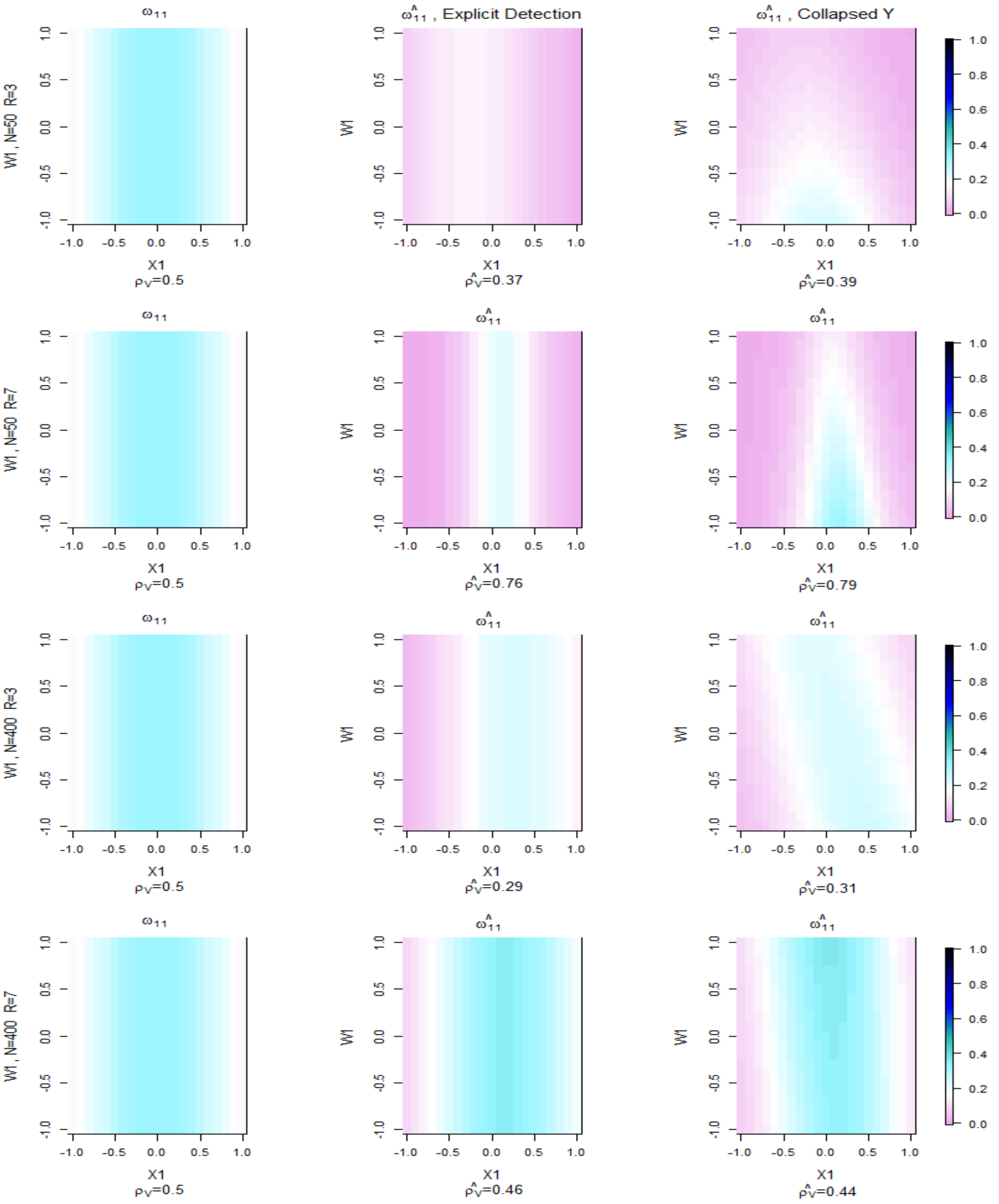


Figure 9: *Comparison of true value of $\omega_{11}$ (left) with mean estimated values $\hat{\omega_{11}}$ for model explicitly accounting for detection (middle) and model using collapsed data (right). Each plot shows $\omega_{11}$ (colour coded) versus covariates $X_1$ and $W_1$. Each row shows the results for different combinations of number of sites (N) and survey replications (R).*

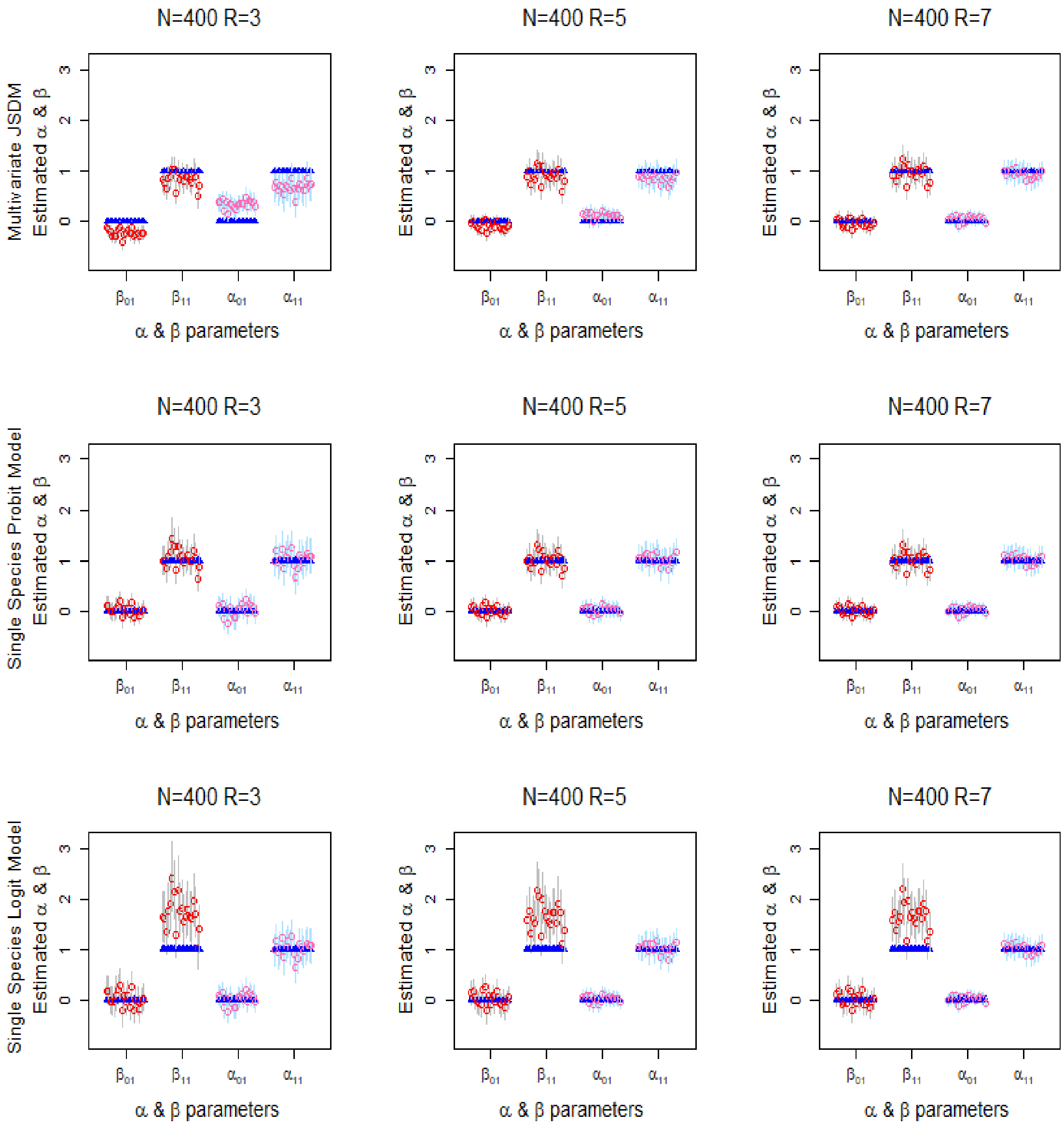


Figure 10: Impact of survey replications $R$ on estimates of occupancy and detection parameters. The top row shows the multivariate JSDM estimates, the middle row show single species logistic estimates, and the last row shows single species probit estimates. Panels show parameter estimates for $R$ increasing from left (3) to right (7) for $N = 400$. Each panel shows 20 estimates for each $\beta$ and $\alpha$ parameter as follows: true value (blue triangle), mean estimate (red circle), 95% credible intervals (grey lines) for occupancy parameters; and mean estimate (pink) and 95% and credible interval (light blue) for detection parameters.

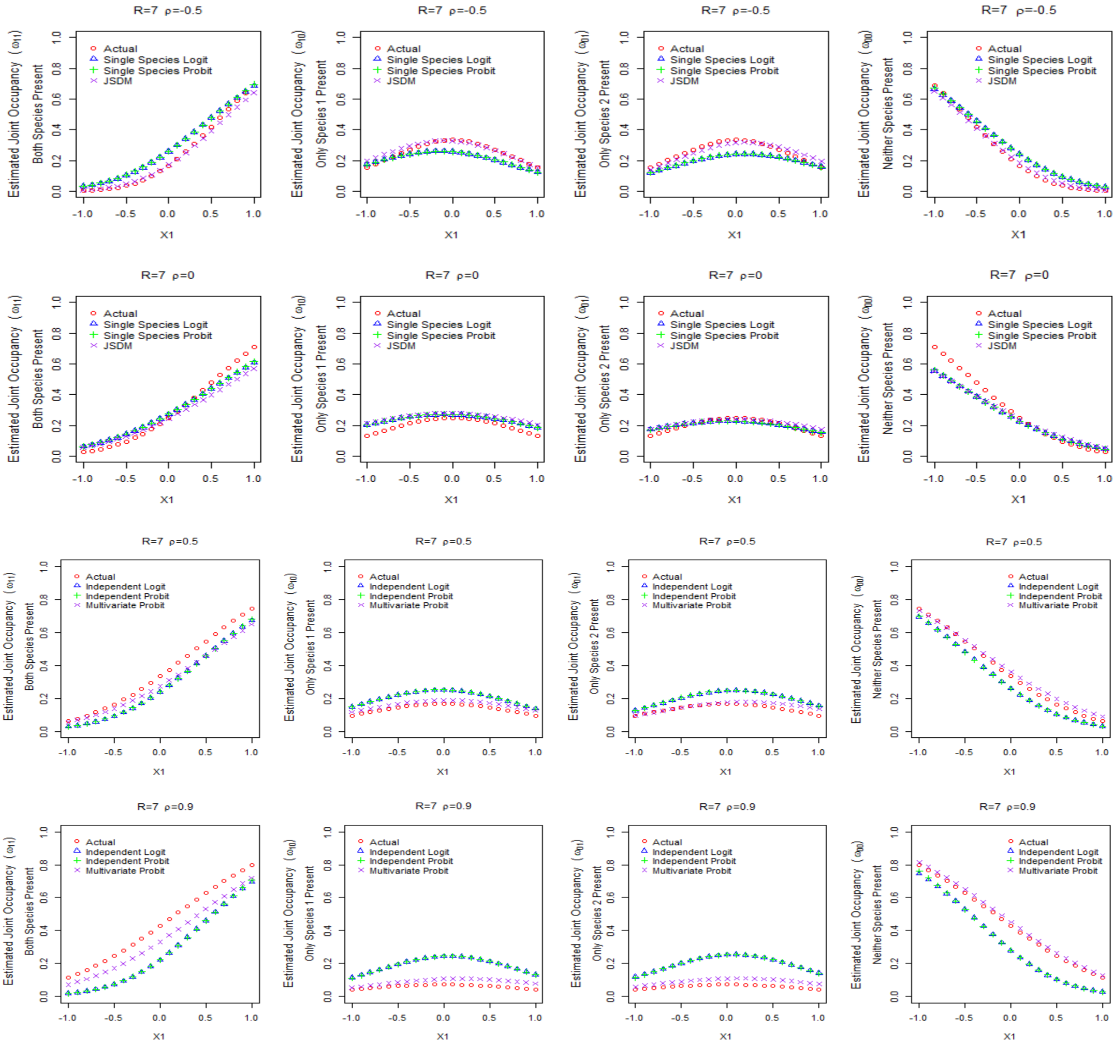


Figure 11: Estimates of $\Omega$ the joint occupancy for covariate case where both species have a positive relation to the occupancy covariate $X_1$. Columns left to right shows estimates of $\omega_{11}$ (probability both species present),$\omega_{01}$,$\omega_{10}$ and ,$\omega_{00}$ (probability neither species present) respectively. Each row shows outcomes as intrinsic correlation $\rho_V$ changes $\rho_V \in \{-0.5, 0, 0.5, 0.9\}$. All estimates are shown for 400 sites and 3 replications. Each panel the joint occupancy probability versus the occupancy covariate $X_1$ for the simulated data (actual - red circles), and each of the models: single species logit (blue triangles), single species probit (green +), and multivariate probit (purple x).

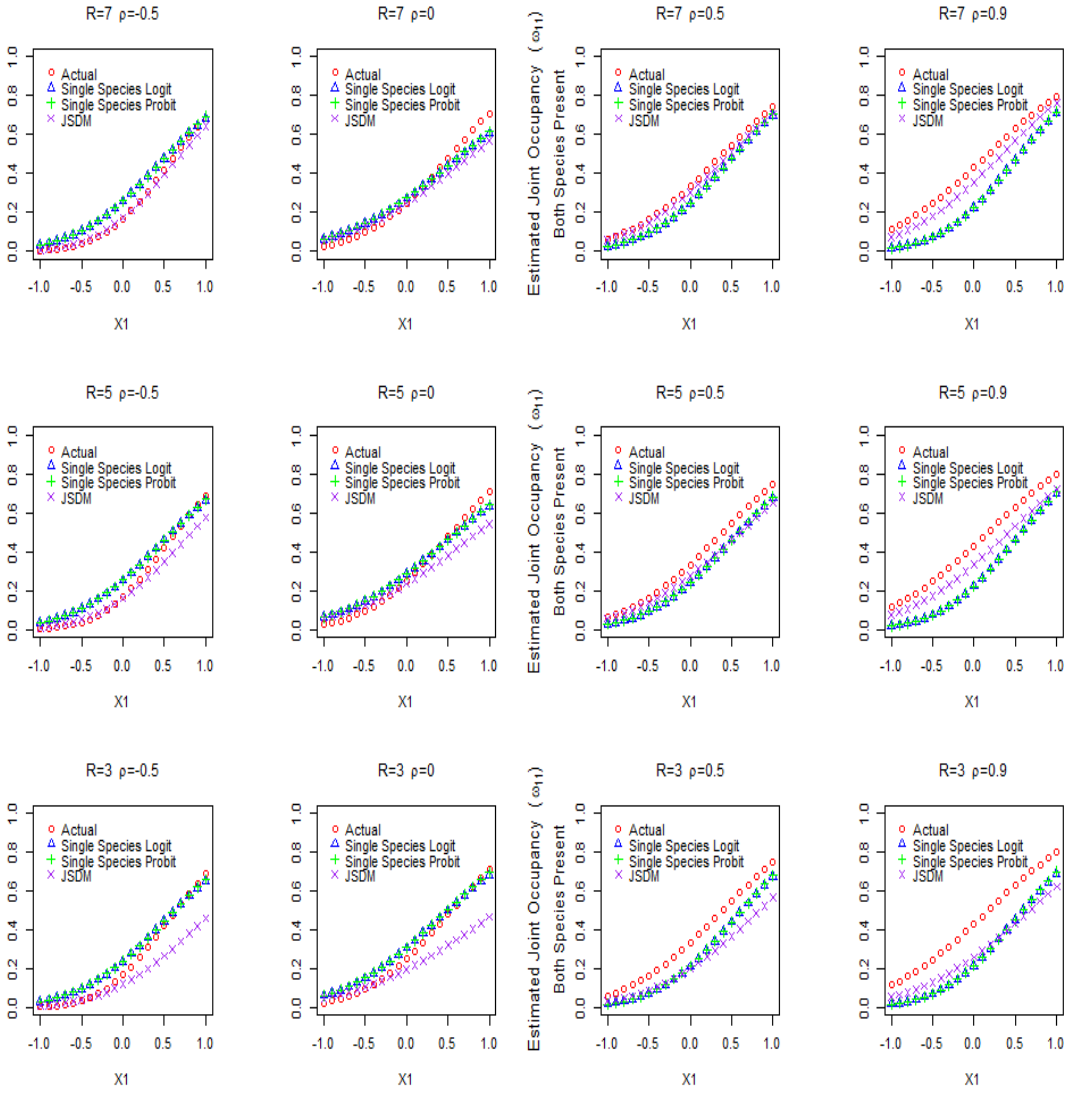


Figure 12: Estimates of $\omega_{11}$, the joint probability both species present, for covariate case where both species have a positive relation to the occupancy covariate $X_1$. Columns left to right shows estimates of $\omega_{11}$ for increasing intrinsic correlation $\rho_V$, where $\rho_V \in \{-0.5, 0, 0.5, 0.9\}$. Each row shows results as number of replications R changes from 7 (top) to 3 (bottom). All estimates are shown for 400 sites. Each panel the joint occupancy probability versus the occupancy covariate $X_1$ for the simulated data (actual - red circles), and each of the models: single species logit (blue triangles), single species probit (green +), and multivariate probit (purple x).

low numbers of replication, with superior estimates for 7 replicates, and bias apparent for other estimates. The slope parameters for the logit model have a much larger magnitude than the other two models, due to the the different link function. The single species probit model mean parameter estimates are generally close to the true value. This is a property of the underlying bivariate normal distribution, where the mean of the marginal distribution is unchanged from the mean over the joint distribution, i.e. if $Z = interval Z_1, Z_2$ have a bivariate normal distribution $Z \sim N([\mu_1, \mu_2], \Sigma)$, then the univariate marginal distribution of $Z_1$ and $Z_2$ are $Z_1 \sim N(\mu_1, \sigma_1 = \sqrt{\Sigma_{11}})$ and $Z_2 \sim N(\mu_2, \sigma_2 = \sqrt{\Sigma_{22}})$. Also errors are small, accuracy improving only slightly as replications increase from 3 to 5

Figures 11 show estimates of the joint distribution probabilities between two species. For two species there are four joint probabilities: $\omega_{11}$ the probability both species are present, $\omega_{10}$ the probability only the first species is present, $\omega_{01}$ the probability only the second species is present, and $\omega_{00}$ the probability neither species is present. These joint probabilities all vary with the occupancy covariate $X_1$. Joint probabilities are calculated from estimated mean values of $\beta$ occupancy parameters and either estimated mean value of $\rho_V$ for the multivariate JSDM or assuming the $\rho_V$ is 0 for other models. These figures show the outcome when all models have their best estimates, with 400 sites and 7 replications. Here the Multivariate JSDM (purple x) generally gives the best response, and most closely follows the line of the actual data (red circles). When species actually are independent (simulated $\rho_V$ is 0), the multivariate JSDM and the single species probit model tend to coincide. As intrinsic correlation increases, the improved estimates become more apparent. The impact of the correlation on estimates of joint probabilities becomes more apparent as the relative environmental component of the estimate decreases, that is, more apparent close to $X_1$ is 0 and less apparent as $X_1$ increases. The single species probit model tends to underestimate the probability that species occur together across most ranges, but over estimate single species occurrence. Calculations for the two single species models (logit and probit) tend to be similar and follow the same curve.

When analysis conditions are not ideal however, the errors in estimates of the $\beta$ occupancy parameters can impact the estimates of joint occupancy. Figure 12 shows how results may change as the number of replications changes. Estimates for 7 replications tend to be very good for the multivariate JSDM compare to the other models. For 5 replications, the multivariate JSDM tends to do better, but not for all ranges of the occupancy covariate: as the covariate value increases the multivariate JSDM joint probability estimate can become less accurate. Finally for 3 replications, the single species probit model does better for a number of $\rho_V$ cases. This is probably due to the larger errors (bias) in estimates of $\beta$ and $\alpha$ parameters for the multivariate JSDM outweighing the utility of the correlation estimate when calculating the joint distribution compared to the single species probit model. Calculations from the two single species models (logit and probit) tend to be very similar across all survey replications.

The result that single species probit models may estimate $\beta$ occupancy parameters and $\alpha$ detection parameters for low numbers of replications could suggest that a sequential Bayesian approach might be able to be employed to estimate correlation more precisely [5]. In such an approach for an initial set of data, a univariate probit could be used to estimate the posterior distribution of the $\beta$ and $\alpha$ parameters. These posterior distributions could then be used in turn as the the prior distributions of these parameters for a subsequent analysis using a secondary data set. The initial data set and secondary datasets could be random subsets of the overall data set.
