## Supplementary material for "Effectiveness of Joint Species Distribution Models in the Presence of Imperfect Detection": Further Case Study Results

### Effectiveness of JSDMs Allowing for Detection Supplement 4: Sensitivity Analysis for Hyper-Priors for Hiearchical Model

Stephanie Hogg, Yan Wang and Lewi Stone

School of Science, RMIT, 124 LaTrobe St, Melbourne, Victoria, 3000, Australia

June 2020

#### 1 Introduction

This supplement provides results for a sensitivity analysis of Bayesian priors for a latent, multivariate probit regression model as used in the paper *Analysis of JSDMs Subject to Imperfect Detection.* It focuses on priors for the intrinsic correlation between species, modelled by the $\Sigma$ correlation matrix.

#### 2 Priors for Intrinsic Covariance and Correlation

A latent, multivariate probit model for a Joint Species Distribution Model was originally proposed by (Pollock *et al.* 2014). This model used a Wishart prior with degrees of freedom *df* set by the number of species $M + 1$. The literature generally agrees that for covariance priors, a Wishart prior with this degree of freedom is generally uninformative (Hurtado Rúa *et al.* 2015).

In our paper, the multivariate probit model was extended to explicitly model detection using an independent Bernoulli component for each species. This extended model shows a bias in estimates of correlation, with the magnitude of correlations is often underestimated when probability of detection is low and number of replications are few.

To determine if result is influenced by the prior, a sensitivity analysis was conducted, comparing the correlation estimates from the wishart prior with the correlation estimates from an alternative approach which uses Multivaraite normal prior $\sim MVN(\mu, \Sigma$, where $\mu$ is the mean and $\Sigma$ is the covariance matrix. Using a Cholesky decomposition of the covariance matrix, $\Sigma$ can be defined in terms of $\mathbf{\Phi}$ and $\mathbf{\Gamma}$, where $\Sigma = \mathbf{\Phi\Gamma\Gamma^T\Phi^T}$. $\mathbf{\Phi} = diag(\phi_{11}, \phi_{22}, \ldots, \phi_{MM})$ and $\mathbf{\Gamma}$, a lower triangual matrix with elements $\gamma_{ij}$, can then be estimated using a series of uniform priors (Demirhan & Hamurkaroglu 2008). These priors can be set with prior knowledge of correlation and tuned by the researches confidence in these estimates.

The priors for each $\phi_{ii}$ is $Uniform(0, h)$, where $h$ sets the confidence of the Correlation estimates: if $h$ is close to 0, then the estimate is considered very uncertain; if $h$ is far from 0, then the estimates are considered quite certain(Demirhan & Hamurkaroglu 2008). For this analysis, values are set a $h = 0.1$ (not confident) and $h = 0.9$ (very confidence) of the estimate.

Priors for each non-zero $\gamma_{ij}$ are also uniform based on an interval which relates to a prior estimate of the correlation. For our data, the prior correlation information is based on the naive correlation found between observations for each species. These value give a rough estimate of the intrinsic correlation, but they are impacted by environmental factors and imperfect detection.

Data was simulated for 2 species over 200 sites for given values of probility of occupancy and detection for correlation values ($rho \in \{-0.75, -0.5, -0.25, 0, 0.25, 0.5, 0.75\}$).

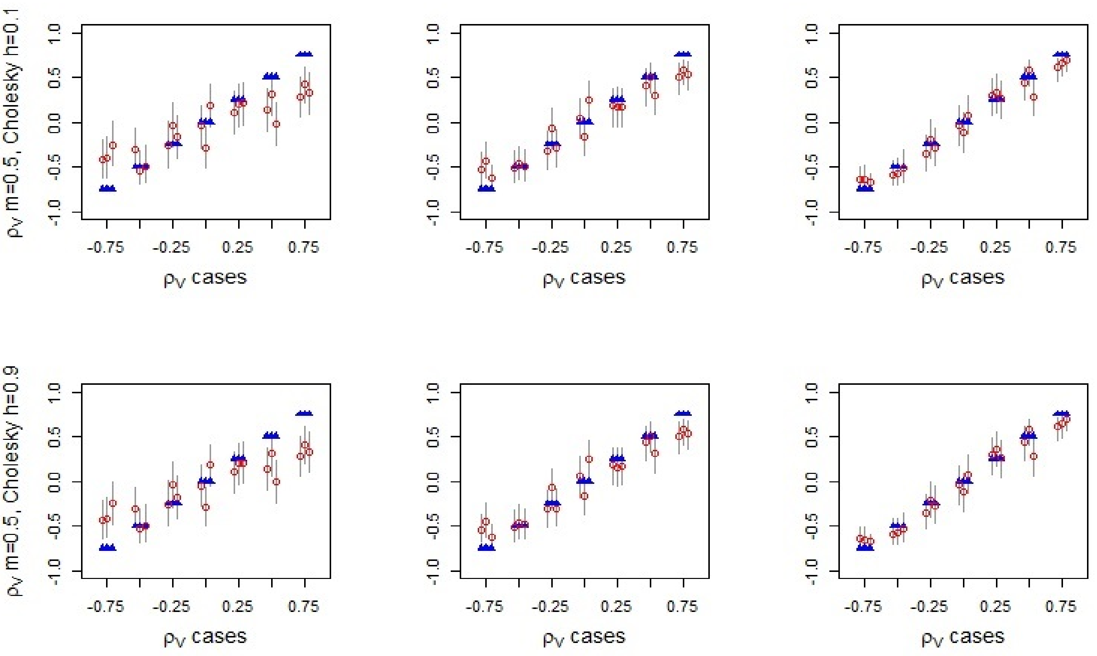


Figure 1: Estimates of correlation $\rho$ for tuneable Cholesky decomposition approach for 2 species and probability of occupancy of 0.5. Each row shows the results for different model types, with $h = 0.1$ model allowing for little confidence in intial estimates and $h = 0.9$ allowing for high confidence in initial estimates. Each panel shows 3 estimates of rho for a series of actual $\rho$ values. Probability of detection $p$ doesn't vary within a panel, but increases across each row.

Figure 1 shows the estimated correlation using the two different h values when 3 observations for each site. We can see these follow a very similar pattern to previous estimates using Wishart correlation with degree of freedom of 3. This suggests that these results are not a result of the priors.

### 3 Hyper Priors for Occupancy and Detection

Normal priors were used for occupancy slope $\beta$ and intercept parameters, i.e. $\beta_{kp} \sim N(\mu_p^{occ}, {\sigma_p^{occ}}^2)$, where $\beta_{kp}$ the parameter for covariate $X_p$ and species $k$. This is a hierarchical model, so hyper priors are used to describe these distributions, with the mean occupancy hyper prior $\mu_p^{occ} \sim Normal(0, \sigma_{occ})$ and the occupancy standard deviation hyper prior $\sigma_p^{occ} \sim Uniform(0, max_{occ})$. A sensitivity analysis was carried out for a number of hyper prior combinations using simulated data. For the mean occupancy hyper-prior, the standard deviation varied with $\sigma_{occ} \in \{4, 8, 10, 100\}$; while for the occupancy standard deviation hyper prior the maximum value varied with $max_{occ} \in \{4, 8, 10, 100\}$. These wide ranges for the standard deviation were chosen based on the knowledge of the restricted (-1,1) range of the simulation parameters.

Normal priors were also used for detection slope $\alpha$ and intercept parameters, i.e. $\alpha_{kp} \sim N(\mu_p^{det}, {\sigma_p^{det}}^2)$, where $\alpha_{kp}$ the parameter for covariate $W_p$ and species $k$. Again, normal and uniform distributions were used as the hyper priors for $\mu_p^{det}$ ($\mu_p^{det} \sim Normal(0, \sigma_{det})$)

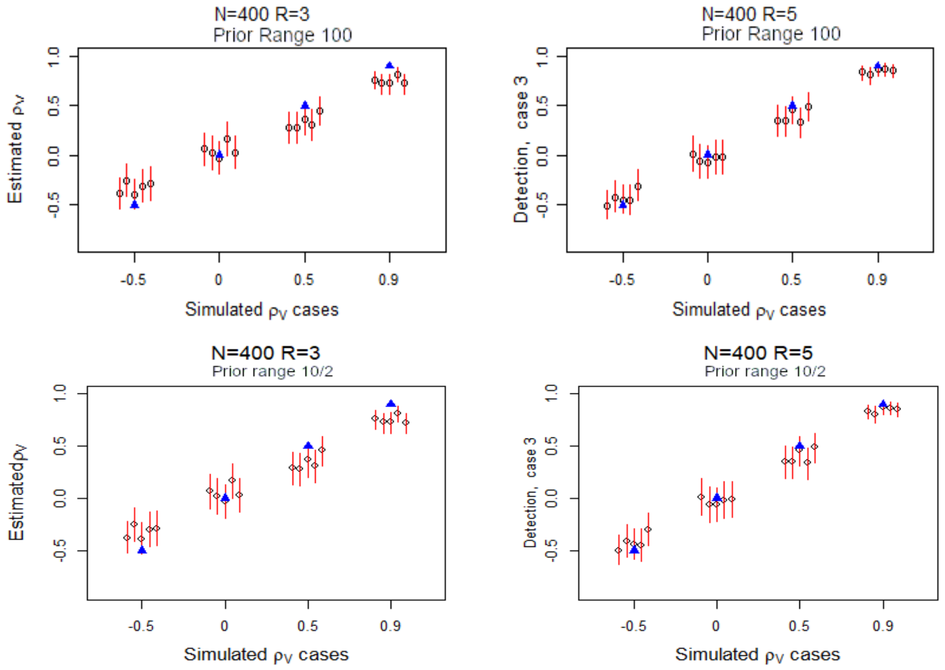


Figure 2: Estimates of correlation $\rho$ for different combinations of $\beta$ occupancy hyper priors and $\alpha$ detection parameter hyper priors. Each row shows the results for different combinations of hyper priors. Each panel shows 5 estimates of rho for a series of $\rho$ simulation cases.

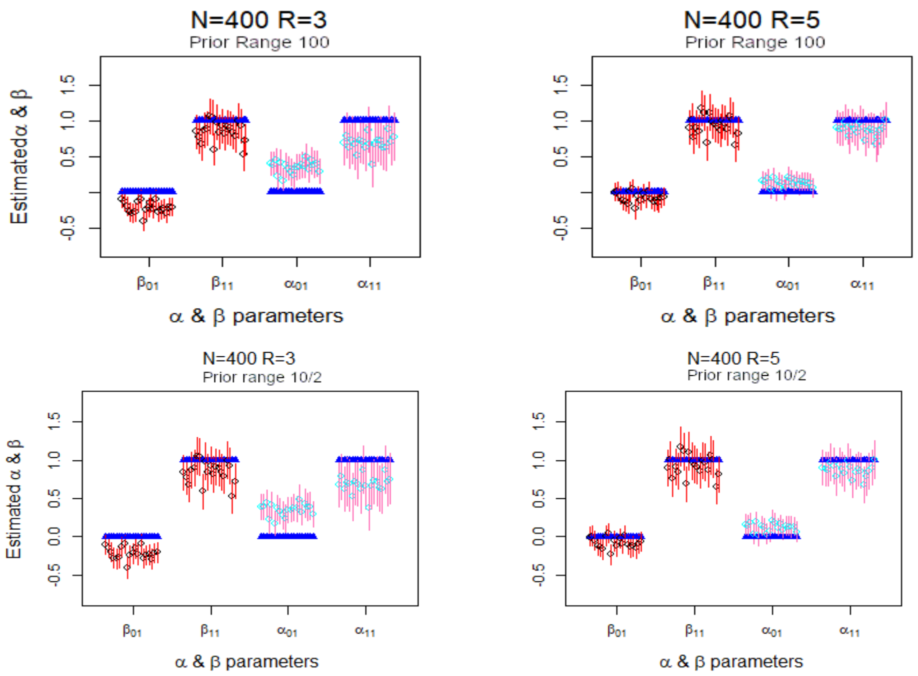


Figure 3: Estimates of occupancy $\beta$ parameters and detection $\alpha$ parameters for different combinations of $\beta$ occupancy hyper priors and $\alpha$ detection parameter hyper priors. Each row shows the results for different combinations of hyper priors. Each panel shows 20 estimates of rho for a one combination of $\alpha$ and $\beta$ parameters, with 3 and 5 survey replications shown.

and $\sigma_p^{det}$ ($\sigma_p^{det} \sim Uniform(0, max_{det})$). Sensitivity analysis of the hyper priors on the detection probability was also conducted, where $\sigma_{det}$ and $max_{det} \in \{100, 6.25, 4, 2\}$.

Analysis showed the varying occupancy and detection hyper prior values did not affect estimates of either detection or occupancy parameters, or estimates of intrinsic correlation between species. Therefore we concluded parameter estimates are not influenced by the choice of prior values for these simulations. While traditionally, some of these priors' values would be considered weakly informative, these results are in line with Lemoine (2019). So all prior values for these simulations may be considered non-informative given the restricted (-1,1) range of the simulation parameters. In later simulations, occupancy hyper priors used $max_{occ} = 4$ and detection hyper priors used $max_{det} = 2$.

#### References

Demirhan, H. & Hamurkaroglu, C. (2008) Bayesian estimation of log odds ratios from r x c and 2 x 2 x k contingency tables. *Statistica Neerlandica*, **62**, 405–424.

Hurtado Rúa, S.M., Mazumdar, M. & Strawderman, R.L. (2015) The choice of prior distribution for a covariance matrix in multivariate meta-analysis: a simulation study. *Statistics in medicine*, **34**, 4083–4104.

Lemoine, N.P. (2019) Moving beyond noninformative priors: why and how to choose weakly informative priors in bayesian analyses. *Oikos*, **128**, 912–928.

Pollock, L.J., Tingley, R., Morris, W.K., Golding, N., O'Hara, R.B., Parris, K.M., Vesk, P.A. & McCarthy, M.A. (2014) Understanding co-occurrence by modelling species simultaneously with a joint species distribution model (jsdm). *Methods in Ecology and Evolution*, **5**, 397–406.
