## Supplementary material for "Effectiveness of Joint Species Distribution Models in the Presence of Imperfect Detection": Sensitivity Analysis

### Effectiveness of JSDMs Allowing for Detection Supplement 3: Victorian Central Highland Owl and Glider Study

[1]School of Science, RMIT, 124 LaTrobe St, Melbourne, Victoria, 3000, Australia

June 2019

#### 1 Study Outline

This document provides supplementary information on the *Victorian Central Highland Owl and Glider Study.* This study is based on surveys conducted in 2014 by the former Victoria Department of Environment and Primary Industry (now the Department of Environment, Land, Water and Planning (DELWP)). This research was in response to the Victorian government's Timber Industry Action Plan (2011) to support the development of a new land management approach [1]. Target species and common abbreviations used are listed below:

Table 1: *Abbreviations for Owl and Glider Data*

| Species Code | Common Name | Official Name |
|---|---|---|
| PO | Powerful Owl | *Ninox strenua* |
| SO | Sooty Owl | *Tyto tenebricosa tenebricosa* |
| GG | Greater Glider | *Petauroides volans* |
| YB | Yellow Bellied Glider | *Petaurus australis* |
| MO | Masked Owl | *Tyto novaehollandiae novaehollandiae* |

#### 2 Parameter Estimates

Parameters for the occupancy and detection of each species were estimated using the model explicitly allowing for detection. Table 2 shows the occupancy parameters - mean values are shown on the top line with the 5% credible interval range shown below in brackets. Credible intervals can be used to deem parameters significant or not. The credible interval is the range between the 2.5% and 97.5% percentiles of all estimates. Here the elevation covariate has a significant parameter for Powerful owls, Sooty Owls and Yellow Bellied (YB) gliders. The greater glider has significant parameters for the TPI (topographic position index) and NVDI (a vegetation index based on satellite spectral analysis).

Table 3 shows the detection parameters, again with mean values on top and 5% credible interval range shown below in brackets. Here the wind speed is a significant for the detection of both Powerful and Sooty Owls. The day of the year (indicator variable distinguishing winter and Autumn observations) is not significant at all. There are no significant detection variables for the glider species.

Table 2: *Occupancy Parameter Estimates for Owl and Glider Data using model explicitly allowing for detection: Mean estimated values top with 5% critical interval values below.*

| Covariate | Parameter | Powerful Owl | Sooty Owl | Greater Glider | YB Glider |
|---|---|---|---|---|---|
| Intercept | $\beta_0$ | -1.012*<br>(-1.298,-0.749) | -0.813*<br>(-1.06,-0.574) | -1.125*<br>(-1.422,-0.843) | -0.969*<br>(-1.264,-0.703) |
| Summer Rain | $\beta_1$ | -0.166<br>(-0.571,0.21) | 0.258<br>(-0.091,0.663) | 0.054<br>(-0.374,0.475) | 0.388<br>(-0.023,0.883) |
| Elevation | $\beta_2$ | -0.441*<br>(-0.798,-0.149) | -0.327*<br>(-0.616,-0.083) | -0.196<br>(-0.491,0.133) | -0.335*<br>(-0.62,-0.082) |
| Wetness index | $\beta_3$ | -0.181<br>(-0.467,0.057) | -0.115<br>(-0.335,0.098) | -0.152<br>(-0.412,0.096) | -0.111<br>(-0.341,0.132) |
| TPI | $\beta_4$ | 0.097<br>(-0.172,0.39) | 0.008<br>(-0.216,0.252) | -0.284*<br>(-0.597,-0.001) | -0.152<br>(-0.417,0.088) |
| NVDI (summer) | $\beta_5$ | 0.223<br>(-0.099,0.579) | -0.114<br>(-0.448,0.196) | 0.439*<br>(0.081,0.858) | 0.244<br>(-0.082,0.603) |

Table 3: *Detection Parameter Estimates for Owl and Glider Data using Explicit Model: Mean estimated values top with 5% critical interval values below.*

| Covariate | Parameter | Powerful Owl | Sooty Owl | Greater Glider | YB Glider |
|---|---|---|---|---|---|
| Intercept | $\alpha_0$ | 0.669*<br>(0.108, 1.205) | 0.630*<br>(0.056, 1.154) | 0.799*<br>(0.279, 1.417) | 0.766*<br>(0.291, 1.293) |
| Wind | $\alpha_1$ | -0.920*<br>(-1.961, -0.076) | -0.866*<br>(-1.804, -0.059) | -0.502<br>(-1.297, 0.415) | -0.574<br>(-1.393, 0.334) |
| Yday | $\alpha_2$ | -0.162<br>(-0.973, 0.626) | 0.083<br>(-0.64, 0.936) | -0.156<br>(-0.98, 0.640) | -0.305<br>(-1.266, 0.498) |

Table 4: *Occupancy Parameter Estimates for Owl and Glider Data using collapsed data model: Mean estimated values top with 5% critical interval values below.*

| Covariate | Parameter | Powerful Owl | Sooty Owl | Greater Glider | YB Glider |
|---|---|---|---|---|---|
| Intercept | $\beta_0$ | -1.047<br>(-1.498,-0.654) | -0.995<br>(-1.441,-0.6) | -1.226<br>(-1.706,-0.767) | -0.838<br>(-1.199,-0.476) |
| Summer Rain | $\beta_1$ | -0.157<br>(-0.574,0.231) | 0.363<br>(-0.011,0.808) | 0.037<br>(-0.411,0.504) | 0.396<br>(-0.008,0.881) |
| Elevation | $\beta_2$ | -0.454<br>(-0.819,-0.143) | -0.443<br>(-0.811,-0.138) | -0.233<br>(-0.549,0.129) | -0.283<br>(-0.559,-0.012) |
| Wetness index | $\beta_3$ | -0.188<br>(-0.469,0.041) | -0.136<br>(-0.365,0.099) | -0.169<br>(-0.458,0.098) | -0.106<br>(-0.336,0.125) |
| TPI | $\beta_4$ | 0.126<br>(-0.144,0.423) | 0.029<br>(-0.217,0.304) | -0.311<br>(-0.622,-0.008) | -0.145<br>(-0.398,0.094) |
| NVDI (summer) | $\beta_5$ | 0.226<br>(-0.118,0.588) | -0.085<br>(-0.435,0.238) | 0.41<br>(0.028,0.852) | 0.195<br>(-0.132,0.547) |
| Wind Speed | $\beta_6$ | -0.101<br>(-0.749,0.493) | -0.107<br>(-0.683,0.448) | 0.289<br>(-0.392,1.133) | -0.18<br>(-0.793,0.347) |
| Yday | $\beta_7$ | 0.157<br>(-0.415,0.759) | 0.437<br>(-0.15,1.101) | 0.076<br>(-0.517,0.632) | -0.284<br>(-1.02,0.305) |

Table 4 shows the occupancy parameter estimates for the collapsed model. Note that there are two more covariates considered for this model as we have included the two detection parameters (see Table 3) also. For each species, the same covariates are significant as in Table 2. However, none of the detection covariates included in this model are significant in this model.

Table 5: Intrinsic Correlation Estimates for Owl and Glider Data For Models Completely Ignoring Detection. Table shows mean estimated intrinsic correlation values on with 95% credible interval values in brackets below. Results are shown for each distinct pair of species, where second species differs from the first species.

| Species | Sooty Owl | Greater Glider | Yellow Bellied Glider |
|---|---|---|---|
| Powerful Owl | 0.0023<br>(-0.4967, 0.4262) | -0.2806<br>(-0.682, 0.2604) | 0.0649<br>(-0.2974, 0.4857) |
| Sooty Owl | | 0.2357<br>(-0.2096, 0.5885) | -0.0824<br>(-0.4445, 0.3075) |
| Greater Glider | | | -0.3566<br>(-0.7148, 0.1473) |

Table 5 shows intrinsic correlation $\rho_V$ estimates for a model that completely ignores detection. This model has no component explicitly accounting for detection, nor does it use collapsed data. Estimates are determined from a single survey without replication. No correlation estimates are significant when detection is completely ignored. The uncertainty is greater for this model when compared with the model explicitly allowing for detection and the model using collapsed data.

### 3 Estimates ROC-AUC

Table 6: *AUC ROC values for the two models*

| Species | Explicitly Accounting for Detection | Collapsed Data |
|---|---|---|
| Powerful Owl | 0.7694 | 0.7667 |
| Sooty Owl | 0.5822 | 0.6639 |
| Greater Glider | 0.5895 | 0.5153 |
| Yellow Bellied glider | 0.7000 | 0.7214 |

AUC-ROC measures were calculated for each species for both the explicit model accounting for detection and the collapsed data model. Table 6 shows the resulting values. AUC-ROC for Powerful Owl's and Yellow Bellied gliders are very similar between the two models. The collapsed model has a higher ROC-AUC for sooty owls and the explicit model has a better ROC-AUC for Greater Gliders. The AUC-ROC for these latter two species is less than that achieved with the previous species for both models, with the ROC-AUC indicating weak predictive power for sooty Owl's and Greater gliders. Over all, ROC-AUC shows a similar performance for the two models when considered across all species.
