## Supplementary material for "Effectiveness of Joint Species Distribution Models in the Presence of Imperfect Detection": Relations Observed and Actual Data

### Effectiveness of JSDMs Allowing for Detection Supplement 5: Theoretical Relations

School of Science, RMIT, 124 LaTrobe St, Melbourne, Victoria, 3000, Australia

October 2020

#### 1 Relations between Observed Correlation and Actual Occupancy using Simple Model of Imperfect Detection

##### 1.1 Simple model of imperfect detection

$Z_i$ represents the observed occupancy of species $i$ at a given location.
$D_i$ represents the chance of seeing a species $i$ at a given location.
$Y_i$ represents the observed occupancy of species $i$ at a given location.
$Y_i$ can be modelled as follows:

$$Y_k = D_k.Z_k \tag{1}$$

This can be extended to the 2 species $h$ and $i$ by considering $Z = (Z_h, Z_i)$, $Y = (Y_h, Y_i)$, $D = (D_h, D_i)$, where $Z_i, Y_i$, and $D_i$ are represent actual occupancy, observed occupancy and the detection process for species $i$. Let us assume that $Z_h$ and $Z_i$ are dependent with correlation $\rho_Z$, but that the detection variables $D_h$ and $D_i$ are independent. Further, assume that the detection process $D_i$ is independent of actual occupancy $Z_i$. We want to understand how the correlation between $Y_h$ and $Y_i$, $\rho_Y$, relates to $\rho_Z$. The relations shown here extend to multiple species if we assume dependencies between species can be represented by pairwise correlation between the species.

##### 1.2 General Relation between Correlation of Y and Z

From first principals, the correlation and covariance of $Z_h$ and $Z_i$ are:

$$\rho_Z = \frac{cov(Z_h, Z_2)}{\sigma_{z_i}.\sigma_{z_2}}$$
$$cov(Z_h, Z_i) = E[Z_h.Z_i] - E[Z_h].E[Z_i]$$

Similarly the correlation and covariance of $Y_h$ and $Y_i$ are given as:

$$\begin{aligned}\rho_Y &= \frac{cov(Y_h, Y_i)}{\sigma_{Y_h}.\sigma_{Y_i}} \\ &= \frac{cov(D_h.Z_h, D_2.Z_2)}{\sigma_{Y_h}.\sigma_{Y_i}}\end{aligned}$$

$$\begin{aligned}cov(Y_h, Y_i) &= cov(D_1.Z_h, D_2.Z_i) \\ &= E[D_h.Z_h.D_i.Z_i] - E[D_h.Z_h].E[D_i.Z_i]\end{aligned}$$

As $D_i$ independent of each other and $Z_i$. From this is is easy to show that:

$$cov(Y_h, Y_i) = \quad = E[D_h].E[D_i]E[Z_h.Z_i] - E[D_h].E[D_i].E[Z_h].E[Z_i]$$
$$= E[D_h].E[D_i].Cov(Z_h.Z_i)$$

In a detection process a species (occupancy) or individual of a species (abundance) is either detected or not. So we can model this detection process by a Bernoulli distribution with probability of detection $p$ and $E(D) = p$. For multiple species, if the detection process is independent, then for each species i, $D_i$ is a Bernoulli distribution with probability $p_i$. So the observed covariance can be written as:

$$cov(Y_h, Y_i) = p_h.p_i.Cov(Z_h, Z_i)$$

The observed correlation can then be written as:

$$\begin{aligned} \rho_Y &= \frac{p_1.p_2.Cov(Z_1.Z_2)}{\sigma_{Y_1}.\sigma_{Y_2}} \\ &= p_1.p_2.\frac{\sigma_{Z_1}.\sigma_{Z_2}}{\sigma_{Y_1}.\sigma_{Y_2}}\rho_Z \end{aligned} \tag{2}$$

This can be represented as:

$$\begin{aligned} &\rho_{Y_{hi}} = \kappa\rho_{Z_{hi}} \\ &\text{Where } \kappa \text{ is defined as:} \kappa = p_h.p_i.\frac{\sigma_{Z_h}.\sigma_{Z_i}}{\sigma_{Y_h}.\sigma_{y_i}} \end{aligned} \tag{3}$$

##### Relation between Variance of Y and Z and $\kappa$

The standard deveviation for the true occupancy of any species can be defined as:

$$\sigma^2_{Z_h} = Var[Z_h] = E(Z_h^2) - E(Z_h)^2$$

We can then determine the standard deviation of the observed occupancy is:

$$\begin{aligned} \sigma^2_{Y_h} &= Var[Y_h] \\ &= E(Y_h^2) - E(Y_h)^2 \\ &= p_h(E(Z_h^2) - p_h.E(Z_h)^2) \end{aligned}$$

This can now be substituded into our previsou defintion of $\kappa$ from Equation 3:

$$\kappa = p_h.p_i.\sqrt[2]{\frac{(E(Z_h^2) - E(Z_h)^2)(E(Z_i^2) - E(Z_i)^2)}{p_h(E(Z_h^2) - p_h.E(Z_h)^2)p_i(E(Z_i^2) - p_h.E(Z_i)^2)}}$$

Simplifying for $p_h$ and $p_i$ and substituting $\mu_{Zh} = E(Z_i)$, we can write this as:

$$\kappa = \sqrt[2]{\frac{p_h p_i(E(Z_h^2) - \mu^2_{Zh})(E(Z_i^2) - \mu^2_{Zi})}{(E(Z_h^2) - p_h\mu^2_{Zh})(E(Z_i^2) - p_h\mu^2_{Zi})}}$$

#### 1.3 Relation between Correlation of Y and Z assuming Bivariate Occupancy

Now we assume that occupancy for the two species is bivariate: that is $Z_h$ and $Z_i$ are bivariate (0 and 1 only). For bivariate random variables $E(Z_h^2) = E(Z_h) = \mu_{Zh}$. The expression for $\kappa$ can be simplified to:

$$\begin{aligned}\kappa &= \sqrt[2]{\frac{p_h p_i(\mu_{Zh} - \mu_{Zh}^2)(\mu_{Zi} - \mu_{Zi}^2)}{(\mu_{Zh} - p_h\mu_{Zh}^2)(\mu_{Zi} - p_h\mu_{Zi}^2)}} \\ &= \sqrt[2]{\frac{p_h p_i(1 - \mu_{Zh})(1 - \mu_{Zi})}{(1 - p_h\mu_{Zh})(1 - p_h\mu_{Zi})}}\end{aligned} \tag{4}$$

**Assuming Bernoulli Distribution of True Occupanc**

If we assume that the distributions of species i and h are Bernoulli with marginal probabilities of occupancy $m_i$ and $m_h$ respectively, then $\mu_{Zi} = E(Z_i) = m_i$ and similarly $\mu_{Zh} = E(Z_i) = m_h$. We can then substitute these into the above equation 4.

$$\kappa = \sqrt[2]{\frac{p_h p_i(1 - m_h)(1 - m_i)}{(1 - p_h m_h)(1 - p_h m_i)}} \tag{5}$$

### 2 Relation of observed occupancy for collapsed data and multiple survey replications.

Show the marginal probability of a species not being observed across multiple survey replications is the same as the marginal probability of a species not being observed in collapsed data:

#### 2.1 Relation if detection is the same for all observations at the same site.

labelSE:ss:ProbObservedOccupancy:Const

*Using the following notation:*
For raw survey data:
$p_{ijk}$ is the conditional probability that species $k$ will be detected at site $i$ during survey $j$, given species $k$ occupies the site; $\mu_{ik}$ is the marginal, mean latent occupancy for species k at site i;
$m_{ik}$ is the marginal probability of occupancy for species k at site i underlying true occupancy data;
We assume that $m_{ik} = \phi(\mu_{ik})$
$p_{ijk}$ is the conditional probability of detection of species $k$ t site $i$ during any survey j given the species actually occupies the site.

For collapsed data:
$Y_{ik}^*$ is the collapsed observation data across R survey at site $i$ for speicies $k$;
$p_{ik}^*$ is is the effective conditional probability of detection for species k at site i for collapsed data for species k is detected at site $i$, given the species is present at the site.

We want to show that the probaility of the species not being detected for the collapsed data is the same as sum of probabilty of not detecting the species across all observations, i.e.: $P(Y^*_{ik} = 0) = P(\Sigma^R_{j=1} Y_{ijk} = 0)$

To start, lets consider the effective conditional probability of detection for species $k$:

$$\begin{aligned}
p^*_{ik} &= P(Y^*_{ik} = 1 \| Z_{ik} = 1) = 1 - P(Y^*_{ik} = 0 | Z_{ik} = 1) \\
&= 1 - P(Y_{ik1} = 0, Y_{ik2} = 0, \ldots, Y_{ikR} = 0 \| Z_{ik} = 1) \\
&= 1 - (1 - p_{ik})^R && (6)
\end{aligned}$$

assuming $p_{ijk}$ is constant across surveys, i.e. $p_{ijk} = p_{ik}$. Now determine the unconditional, marginal probability that species k is detected for collapsed data:

$$P(Y^*_{ik} = 1) = P(Y^*_{ik} = 1 \| Z_{ik} = 1) P(Z_{ik} = 1) + P(Y^*_{ik} = 1 \| Z_{ik} = 0) P(Z_{ik} = 0) \quad (7)$$

Now we know that:

$$\begin{aligned}
P(Y^*_{ik} = 1 \| Z_{ik} = 0) &= 0 \\
P(Z_{ik} = 1) &= m_{ik} \\
P(Z_{ik} = 0) &= 1 - m_{ik} \\
P(Y^*_{ik} = 1 \| Z_{ik} = 1) &= p^*_{ik} = 1 - (1 - pik)^R)
\end{aligned}$$

Hence:

$$P(Y^*_{ik} = 1 - m_{ik}(1 - p)ik^R \quad (8)$$

And futher,

$$\begin{aligned}
P(Y^*_{ik} = 0) &= P(Y^*_{ik} = 0 \| Z_{ik} = 1) P(Z_{ik} = 1) + P(Y^*_{ik} = 0 \| Z_{ik} = 0) P(Z_{ik} = 0) \\
&= (1 - p^*_{ik}) m_{ik} + 1(1 - m_{ik}) \\
&= 1 - m_{ik} + m_{ik}(1 - p_{ik})^R && (9)
\end{aligned}$$

Now lets consider the model with explicit detection and the probabilities of observing a species for any single observation.

$$\begin{aligned}
P(Y_{ijk} = 1) &= P(Y_{ijk} = 1 \| Z_{ik} = 1) P(Z_{ik} = 1) + P(Y_{ijk} = 1 \| Z_{ik} = 0) P(Z_{ik} = 0) \\
&= p_{ik} m_{ik} + 0(1 - m_{ik} = p_{ik} m_{ik} \\
P(Y_{ijk} = 0) &= 1 - P(Y_{ijk}) = 1 - p_{ik} m_{ik} && (10)
\end{aligned}$$

The unconditional probability that all observations are 0, assuming $p_{ik}$ does not vary between observations.

$$\begin{aligned}
P(\Sigma^R_{j=1} Y_{ijk} = 0) &= P(Y_{i1k} \text{ and } Y_{i2k} \text{ and } \ldots Y_{iRk} = 0) \\
&= P(Y_{i1k} \text{ and } Y_{i2k} \text{ and } \ldots Y_{iRk} = 0 \| Z_{ik} = 0) P(Z_{ik} = 0) \\
&\quad + P(Y_{i1k} \text{ and } Y_{i2k} \text{ and } \ldots Y_{iRk} = 0 \| Z_{ik} = 1) P(Z_{ik} = 1) && (11)
\end{aligned}$$

Now:

$$\begin{aligned}
P(Y_{i1k} \text{ and } Y_{i2k} \text{ and } \ldots Y_{iRk} = 0 | Z_{ik=0}) &= 1 \\
P(Z_{ik} = 1) &= m_{ik} \\
P(Z_{ik} = 0) &= 1 - m_{ik} \\
P(Y_{i1k} \text{ and } Y_{i2k} \text{ and } \ldots Y_{iRk} = 0 \| Z_{ik} = 1) &= \Pi^R_{j=1}(1 - p_{ik}) = (1 - p_{ik})^R
\end{aligned}$$

Hence

$$P(\Sigma_{j=1}^{R} Y_{ijk} = 0) = 1 - m_{ik} + m_{ik}(1 - p_{ik})^R \tag{12}$$

Hence we have shown that: $P(Y^*_{ik} = 0) = P(\Sigma_{j=1}^{R} Y_{ijk} = 0)$

#### 2.2 Relation using a binomial approach

This could also be shown using the binomial approach. For Survey data with replications, the probability that the number of observations across all survey replications can be considered a binomial. Because our data is binary, then the conditional probabily the species is detected $n$ times over our $R$ surveys can be considerd a binomial if $p_{ijk}$ doesn't change between surveys i.e. $p_{ijk} = p_{ik} \forall j \in \{1, \ldots, R\}$ and the species occurs at the site. In this case:

$$\begin{aligned} P(\Sigma_{j=1}^{R} Y_{ijk} = n \| Z_{ik} = 1) &= \text{P(n detections of species k in R surveys)} \\ &= {}^{R}C_n p_{ik}^n (1 - p_{ik})^{R-n} \end{aligned}$$

So if $n = 0$, then

$$\begin{aligned} P(\Sigma_{j=1}^{R} Y_{ijk} = 0 \| Z_{ik} = 1) &= \text{P(0 detections of species k in R surveys)} \\ &= {}^{R}C_0 p_{ik}^0 (1 - p_{ik})^{(R-0)} \end{aligned}$$

Now ${}^{R}C_0 = 1$, and $p_{ik}^0 = 1$, so

$$P(\Sigma_{j=1}^{R} Y_{ijk} = 0 \| Z_{ik} = 1) = (1 - p_{ik})^R \tag{13}$$

Now the unconditional proability of observing the species across all survey replication is

$$\begin{aligned} P(\Sigma_{j=1}^{R} Y_{ijk} = 0) = P(\Sigma_{j=1}^{R} Y_{ijk} = 0 | Z_{ik} = 0).P(Z_{ik} = 0) \\ + P(\Sigma_{j=1}^{R} Y_{ijk} = 0 | Z_{ik} = 0).P(Z_{ik} = 1) \end{aligned}$$

As before, $P(\Sigma_{j=1}^{R} Y_{ijk} = 0 | Z_{ik} = 0) = 1$ by definition. The probability the site is actually occupied is $P(Z_{ik} = 1) = m_{ik}$, while the probability it is not occupied is $P(Z_{ik} = 0) = 1 - m_{ik}$. The unconditional probability of observation can then be written as:

$$P(\Sigma_{j=1}^{R} Y_{ijk} = 0) = 1 - m_{ik} + m_{ik}(1 - p_{ik})^R \tag{14}$$

So as before $P(Y^*_{ik} = 0) = P(\Sigma_{j=1}^{R} Y_{ijk} = 0)$

#### 2.3 Relation if detection varies across observations at the same site.

labelSE:ss:ProbObservedOccupancy:Varies

However, if $p_{ik}$ is not constant, then a binomial approach no longer applies but none the less the general point holds as is shown below:

For the collapsed model, the probability $Y^*_{ik} = 0$, the following now hold:

$$\begin{aligned} P(Z_{ik} = 1) &= m_{ik} \\ P(Z_{ik} = 0) &= 1 - m_{ik} \\ P(Y^*_{ik} = 1 \| Z_{ik} = 0) &= 0 \\ P(Y^*_{ik} = 1 \| Z_{ik} = 1) &= 1 - \Pi_{j=1}^{R}(1 - p_{ijk}) \end{aligned}$$

So the unconditional probabilities of detection for collapsed data become:

$$P(Y^*_{ik} = 1) = m_{ik}(1 - \Pi^R_{j=1}(1 - p_{ijk}))$$
$$P(Y^*_{ik} = 0) = 1 - m_{ik}(1 - \Pi^R_{j=1}(1 - p_{ijk})) \quad (15)$$

Similarly, for the model with detection, using the same approach as before:

$$= 1 - m_{ik} + m_{ik}(\Pi^R_{j=1}(1 - p_{ijk}))P(\Sigma^R_{j=1}Y_{ijk} = 0) \quad (16)$$
